# An optimised transformation protocol for *Anthoceros agrestis* and three more hornwort species

**DOI:** 10.1101/2022.08.10.503456

**Authors:** Manuel Waller, Eftychios Frangedakis, Alan Marron, Susanna Sauret-Gueto, Jenna Rever, Cyrus Raja Rubenstein Sabbagh, Julian M. Hibberd, Jim Haseloff, Karen Renzaglia, Péter Szövényi

## Abstract

Land plants comprise two large monophyletic lineages, the vascular plants and the bryophytes, which diverged from their most recent common ancestor approximately 480 million years ago. Of the three lineages of bryophytes, only the mosses and the liverworts are systematically investigated, while the hornworts are understudied. Despite their importance for understanding fundamental questions of land plant evolution, they only recently became amenable to experimental investigation, with *Anthoceros agrestis* being developed as a hornwort model system. Availability of a high quality genome assembly and a recently developed genetic transformation technique makes *A. agrestis* an attractive model species for hornworts. Here we describe an updated and optimised transformation protocol for *A. agrestis* which can be successfully used to genetically modify one more strain of *A. agrestis* and three more hornwort species, *Anthoceros punctatus, Leiosporoceros dussi* and *Phaeoceros carolinianus.* The new transformation method is less laborious, faster and results in the generation of greatly increased numbers of transformants compared to the previous method. We have also developed a new selection marker for transformation. Finally, we report the development of a set of different cellular localisation signal peptides for hornworts providing new tools to better understand hornwort cell biology.

## Introduction

Hornworts, together with the mosses and liverworts, belong to bryophytes, a monophyletic group sister to all other land plants (tracheophytes). Bryophytes are crucial for revealing the nature of the land plant common ancestor and improving our understanding of fundamental land plant evolutionary innovations. Until recently, model organisms were only available for the mosses and the liverworts, while a tractable model system was lacking for the third group of bryophytes, the hornworts. To further our understanding of hornwort biology, *Anthoceros agrestis* was recently proposed as an experimental model system (Frangedakis, Shimamura, et al. 2021; Szövényi et al. 2015; Gunadi, Li, and Van Eck 2022; Neubauer et al. 2022; Li et al. 2020) with two geographic isolates (Oxford and Bonn) available. Axenic culturing methods for both isolates have been established, their genomes have been sequenced and an *Agrobacterium*-mediated transformation method has been developed (Frangedakis et al., 2021). These technical advances provide us with the tools to unlock fundamental questions of hornwort biology.

While the published transformation method was applied successfully to recover stable transformants for the Oxford isolate of *A. agrestis*, it resulted only in a very low number of stable transformants for the Bonn isolate. The Bonn isolate (Figure 1 A-D) is of particular interest for embryo and sporophyte development studies since, in contrast to the Oxford isolate, sporophytes can be induced en masse under axenic conditions (Supp Fig 1). Sporophytes of hornworts are morphologically and developmentally very different from other bryophytes. Understanding the genetic pathways controlling their development, in comparison to the sporophyte development of mosses and liverworts, is crucial to obtain a complete picture on the evolution of sporophyte body plans in bryophytes and ultimately land plants.

**Figure 1:**
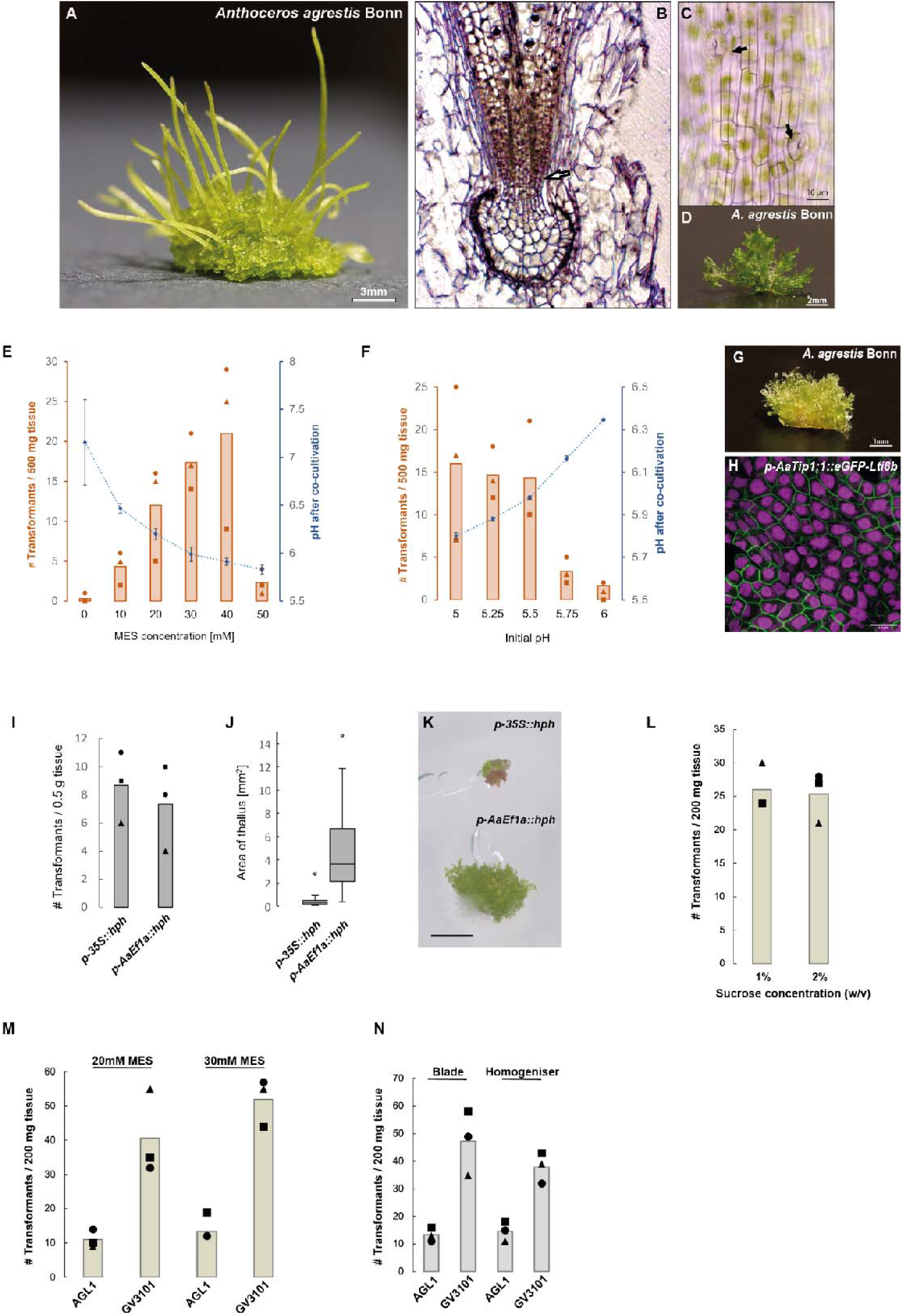
Characteristics of the *A. agrestis* Bonn strain and results of the optimization experiments to improve transformation efficiency. *A-D:* A) *Anthoceros agrestis* Bonn plant bearing sporophytes. Scale bar: 3 mm. B) Longitudinal section of the foot surrounded by the involucre showing the basal meristem (arrow) and differentiating sporogenous tissue. C) Light microscopy image of the sporophyte epidermis showing stomata (see arrows). Scale bar: 10 μm. D) Image of *A. agrestis* Bonn thallus fragment (prior to razor blade or ultra-turrax aided fragmentation) used as starting material for the co-cultivation with *Agrobacterium*. Scale bar: 2 mm. *E-F:* Comparison of the number of transformants (per 500 mg starting tissue) and final pH of the co-cultivation media (after 3 days co-cultivation) for E) different MES concentrations in the co-cultivation media (initial pH set to 5.5) and F) different initial pH values of the co-cultivation media with 40 mM MES. Graphs show values of triplicate experiments (symbols) and their average (bars). Error bars on pH values depicting SDs. *G-H:* G) Transgenic *A. agrestis* Bonn transformed with the *p-AaEf1a::hph - p- AaTip1;1::eGFP-Lti6b* construct. Scale bar: 1 mm. H) Confocal microscopy image of *A. agrestis* Bonn gametophyte transformed with the *p-AaEf1a::hph - p-AaTip1;1::eGFP-Lti6b* construct. Scale bar: 50 μm. Construct maps at Supp Table 1. *I-K:* Effect of the promoter (*AaEf1a* versus the CaMV 35S) driving the hygromycin selection cassette. I) Histogram showing numbers of recovered independent transformant lines expressing GFP. J) Boxplot showing distribution of thallus size between the two constructs (box depicts quartile and median, with whiskers showing max. and min. values inside 1.5x IQR, outliers outside that range depicted as dots. K) Top: Transgenic *A. agrestis* Bonn transformed with the *p-35S::hph - p-35S_s::eGFP-Lti6b* construct, Bottom: Transgenic *A. agrestis* Bonn transformed with the *p-AaEf1a::hph - p-AaTip1;1::eGFP-Lti6b* construct. Scale bar: 1 mm. *L-N:* Comparison of the number of transformants (per 200 mg starting tissue) (after 3 days co-cultivation) for L) different sucrose concentrations in the transformation buffer (initial pH set to 5.5, 20mM MES), M) different *Agrobacterium* strains used for the transformation (tested with *A. agrestis* Bonn), and N) different fragmentation methods (tested with *A. agrestis* Bonn).

**Table 1:** eGFP expression patterns in the stable transformants.

|  | *construct* | *Species/s*<br>*train* | *Number of*<br>*transformants* | *Expression*<br>*in rhizoids* | *Patchy*<br>*expression* | *No*<br>*fluorescence* |
| --- | --- | --- | --- | --- | --- | --- |
|  | *p-AaEf1a::hph - p-AaEF1a::eGFP-Lti6b* | *Oxford* | 211 | 22 | 13 | 10 |
|  | *p-AaEf1a::hph - p-AaEF1a::eGFP-Lti6b* | *Bonn* | 78 | 17 | 8 | 7 |
|  | *p-AaEf1a::hph - p-AaEF1a::eGFP-Lti6b* | *Bonn*<br>*GV3101* | 122 | 22 | 15 | 11 |
|  | *p-AaEf1a::hph - p-AaEF1a::eGFP-Lti6b* | *A.*<br>*punctatus* | 28 | 5 | 4 | 1 |
|  | *p-AaEf1a::hph - p-35S_s::eGFP-Lti6b* | *Bonn* | 8 | 1 | - | 3 |
|  | *p-AaEf1a::hph - p-35S_s::eGFP-Lti6b* | *Oxford* | 146 | 21 | 10 | 7 |
|  | *p-AaEf1a::hph - p-AaTip1;1::eGFP-Lti6b* | *Bonn* | 34 | 10 | 10 | 5 |
|  | *p-AaEf1a::hph - p-AaTip1;1::eGFP-Lti6b* | *Oxford* | 16 | 1 | 4 | 1 |
|  | *p-AaEf1a::hph - p-AaTip1;1::eGFP-Lti6b* | *A.*<br>*punctatus* | 7 | - | - | 2 |

Our goal was to optimise the previously published transformation protocol for the *A. agrestis* Bonn strain and potentially apply it to other hornwort species. To do so, we examined a range of approaches and parameters that were most likely to influence the infection efficiency of *Agrobacterium* and the recovery of stable transformants: 1) the effect of the pH of the co-cultivation media 2) the effect of the promoter driving the selection marker, 3) the sucrose concentration of the co-cultivation solution and 4) the choice of the *Agrobacterium* strain used. We successfully developed a modification of the Frangedakis et al. 2021 protocol, that allowed us to obtain up to 55 stable independent transformant lines from 200 mg of tissue for the *A. agrestis* Bonn isolate and 103 stable transformant lines from 200 mg of tissue for the Oxford isolate We then tested whether the optimised transformation method is applicable to three other hornwort species: *Anthoceros punctatus, Leiosporoceros dussii,* and *Phaeoceros carolinianus. A. punctatus* has long been used as the study system for hornwort associations with cyanobacteria (*Nostoc punctiforme*) (Chatterjee et al. 2022). *Leiosporoceros dussii* is the sister taxon to all hornworts (Duff et al. 2007). Its chloroplasts lack a pyrenoid and one of its key morphological innovations include a unique symbiotic arrangement of endophytic cyanobacteria (Villarreal A and Renzaglia 2006; Villarreal A et al. 2018). *P. carolinianus* represents a commonly found species that can grow relatively easily in laboratory conditions.

Finally, in order to facilitate future studies of hornworts at a cellular level, we developed a set of Loop cloning system/OpenPlant kit compatible tags for localisation at the mitochondria, Golgi, peroxisomes, actin cytoskeleton, chloroplasts and the endoplasmic reticulum (ER) and tested the utility of an additional fluorescent protein and chlorsulfuron as new selection marker for transformation.

In summary, we provide a streamlined transformation protocol which can be used for a series of hornwort species and can potentially be applied to further species of hornworts.

## Results

### Effect of MES concentration on transformation efficiency

It has been reported that a stable pH of 5.5 during co-cultivation leads to significantly increased *Agrobacterium*-infection of *Arabidopsis* cells, likely due to inhibited calcium-mediated defence signalling (Wang et al. 2018). Since this innate immune response is possibly conserved among land plants, the pH during co-cultivation may be a crucial factor for increased *Agrobacterium*-infection rates of hornwort cells. While the published *A. agrestis* transformation method is employing phosphate buffered KNOP medium with a pH of 5.8 for co-cultivation, monitoring pH during co-cultivation of *A. agrestis* with *Agrobacterium*, revealed a pH value increase from an initial value of 5.8 to around 7-8, likely due to *Agrobacterium* growth. A test trial with the *p-AaEf1a::hph - p-AaTip1;1::eGFP-Lti6b* construct (*AaEf1a*: *A. agrestis Elongation Factor 1a, hph:* hygromycin B resistance *phosphotransferase, AaTip1;1: A. agrestis Gamma Tonoplast Intrinsic Protein 1;1,* full list of all construct maps used in this study in Supp Table 1), the initial pH set to 5.5 and the co-cultivation medium supplemented with 10 mM 4-Morpholineethanesulfonic acid (MES) for improved buffering, increased the number of recovered stable transformants for *A. agrestis* Bonn. In this pre-trial we also replaced the use of a homogeniser to fragment the thallus tissue (Supp Fig 2) prior to the co-cultivation with a razor blade, to eliminate the need for special equipment.

To further test the influence of pH range and stability during co-cultivation on transformation efficiency, we added MES in a range of concentrations, 0-50 mM, to the co-cultivation medium and evaluated the effect of different pH values on transformation efficiency. Increasing the concentration of MES in the co-cultivation medium led to a slower rise of pH during co-cultivation and greatly increased the number of transformants (Fig. 1E). However, MES at concentrations higher than 40mM had an apparent toxic effect on *A. agrestis* gametophyte tissue (Supp Fig 3). As indicated by chloroplasts turning brown or grey, plant fragments immediately after co-cultivation looked generally less healthy compared to plants grown under lower than 40mM MES concentrations. When applying a MES concentration of 50 mM, most plant fragments appeared to be dead within 4-5 days after co-cultivation, leading to a decrease in the total amount of recovered transformants. The optimal MES concentration was therefore determined to be 40 mM MES.

### Effect of initial pH of co-cultivation media on transformation efficiency

We also tested the effect of the initial pH of the co-cultivation medium on transformation efficiency. Comparison of different initial pH from 5 to 5.5 showed no significant difference in transformation efficiency, but the number of transformants decreased when the starting pH was set to 5.75 or higher (Fig. 1F-H). In conclusion, the highest number of stable transformants could be recovered with an initial pH between 5-5.5 and 40 mM MES for improved buffering. It should be noted that this is only valid for the specific tissue culture conditions used in our trials, and using tissue thallus fragments that have been propagated for up to 6 weeks after sub-subculturing (Supp Fig 3). Nonempirical observations suggest that thallus tissue from older cultures (2-3 months after subculturing) show a higher sensitivity to MES, with 20 mM MES frequently killing the majority of cells. In this case, higher numbers of recovered stable transformants were usually achieved with only 10 mM MES.

### Effect of promoter on selection and recovery of transformant lines

The promoter used to drive expression of the selection gene hygromycin B resistance-encoding *phosphotransferase* (*hph*) may influence the number of recovered stable transformants. The cauliflower-mosaic virus (CaMV) 35S promoter previously used for this purpose shows stronger expression in differentiated cells of the thallus tissue and weaker expression in new grown parts of the thalli (Frangedakis, Waller, et al. 2021). In contrast, the endogenous *AaEf1a* promoter is more strongly expressed in new grown tissue of the Oxford isolate (Frangedakis, Waller, et al. 2021), which we hypothesise to be more susceptible to *Agrobacterium* infection. Interestingly, the *AaEf1a* promoter shows high activity in protoplasts of *A. agrestis Bonn*, while CaMV 35S activity could not be detected (Neubauer et al. 2022). Thus, we reasoned that driving expression of the *hph* under the *AaEf1a* promoter might improve transformation efficiency.

To test the effect on transformation efficiency when driving the expression of the hygromycin resistance (*hph*) gene with either the *AaEf1a* or the CaMV 35S promoter, we transformed equal amounts of *A. agrestis* Bonn tissue with two different vectors. One contained a *p- 35S::hph* transcription unit and the other a *p-AaEf1a::hph* transcription unit. Both vectors further contained a *p-AaTip1;1::eGFP-Lti6b* transcription unit to allow the use of fluorescence as an additional marker for successful transformation. We then compared the number of isolated eGFP expressing transformants and their growth after a total of seven weeks on selective media. For both constructs, the number of primary transformants (thalli) expressing eGFP in at least some cells of the thallus was comparable (Fig. 1I). However, the majority of the plants carrying the *hph* gene driven by the CaMV 35S promoter showed retarded growth compared with plants carrying the *hph* gene driven by the *AaEf1a* promoter. In particular, after seven weeks of growth, the surface area of thalli of transformants carrying the *p-AaEf1a::hph* transcription unit, was on average ten times that of plants carrying the *p- 35S::hph* transcription unit (Fig. 1J and K) (based on three independent experimental replicates). So while the number of initially isolated transformant lines is not affected by the choice of the promoter driving *hph*, it does have a large effect on the recovery and growth rate of transformed thalli.

### Effect of sucrose concentration of the co-cultivation medium on transformation efficiency

We compared the effect on transformation efficiency of two different concentrations of sucrose, 1% (w/v) and 2% (w/v). We reasoned that lowering the concentration of sucrose in the co-cultivation medium might reduce the chances of *Agrobacterium* overgrowth during co-cultivation that can potentially affect the pH of the medium and transformation efficiency. We infected equal amounts of *A. agrestis* Bonn tissue using co-cultivation media with 20mM MES, the *AGL1 Agrobacterium* harbouring the *p-AaEf1a::hph - p-35S_s::eGFP-Liti6b* construct and with either 1% (w/v) or 2% (w/v) sucrose concentration. Our results suggest that the transformation efficiency is comparable (Fig. 1L).

### Effect of Agrobacterium strain used

Previously, only a very small number of transformants were obtained when the *GV3101 Agrobacterium* strain was used. We tested again the *GV3101 Agrobacterium* strain for its ability to infect and transform *A. agrestis* thallus compared to the *AGL1* strain. We infected equal amounts of *A. agrestis* Bonn and Oxford tissue using co-cultivation media with 20mM MES with *GV3101* or *AGL1 Agrobacterium* harbouring the *p-AaEf1a::hph - p-AaEf1a::eGFP- Liti6b* construct. We repeated the same experiment using co-cultivation media with 20mM and 30mM MES concentration. Our results revealed that when the *GV3101* strain was used the number of transformants was greater (Fig. 1M). This contradicts our previous findings (Frangedakis, Waller, et al. 2021) which could be explained by differences in the quality of *GV3101 Agrobacterium* batches, and/or the difference in pH stability during co-cultivation.

### Comparison of homogenised vs. blade fragmented tissue

To test the effect of tissue fragmentation method on transformation efficiency we infected equal amounts of *A. agrestis* Bonn tissue fragmented using a razor blade or a homogeniser and using co-cultivation media with 30mM MES and the *AGL1* or *GV3101 Agrobacterium* harbouring the *p-AaEf1a::hph - p-AaEf1a::eGFP-Liti6b* construct. Our results show that when using a razor blade the transformation efficiency is slightly higher compared to using a homogenizer (Fig. 1N).

### Comparison of transformation efficiency between the two *A. agrestis isolates* and *A. punctatus*

We then confirmed that the protocol can successfully be used to recover stable transformants for the *A. agrestis* Oxford isolate (Fig. 2A-D).

**Figure 2:**
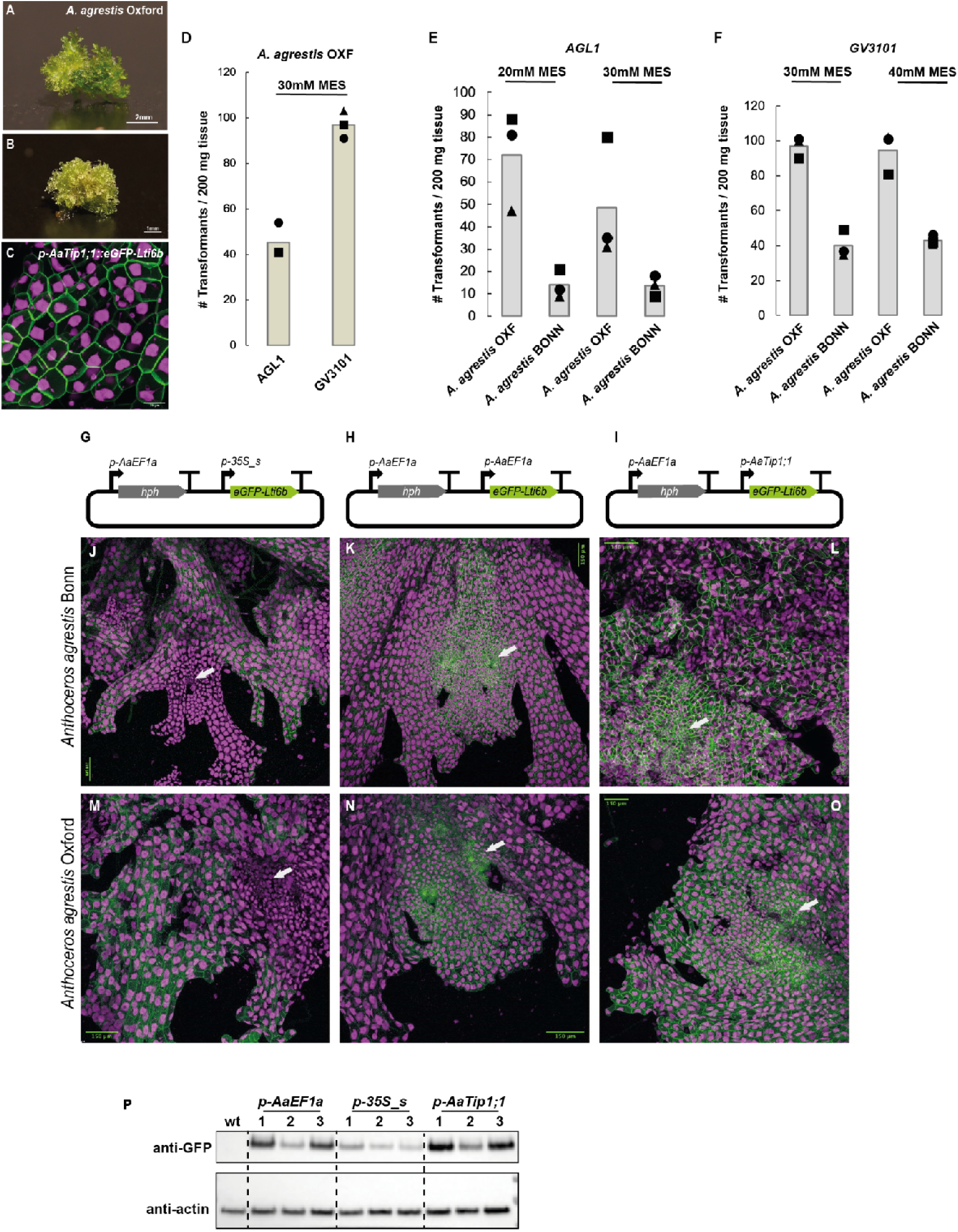
Results of transformation efficiency optimization experiments for the *A. agrestis* isolates, schematic representation of transformation constructs and confocal microscopy images of transgenic *A. agrestis* isolates. A) Image of *A. agrestis* Oxford thallus fragment used as starting material for the co- cultivation with *Agrobacterium*. Scale bar: 2 mm. B) Transgenic *A. agrestis* Oxford gametophyte transformed with the *p-AaEf1a::hph - p-AaTip1;1::eGFP-Lti6b* construct. Scale bar: 1 mm. C) Confocal microscopy image of *A. agrestis* Oxford gametophyte transformed with the *p-AaEf1a::hph - p-AaTip1;1::eGFP-Lti6b* construct. Scale bar: 25 μm. D) Comparison of the number of transformants (per 200 mg starting tissue) (after 3 days co- cultivation) for *A. agrestis* Oxford isolate using two different *Agrobacterium* strains (*AGL1* and *GV3101*). E) Comparison of number of transformants (per 200 mg starting tissue) (after 3 days co-cultivation) for the two *A. agrestis* isolates under two different MES concentrations in the transformation buffer (initial pH set to 5.5) using the *AGL1 Agrobacterium* strain. F) Comparison of number of transformants (per 200 mg starting tissue) (after 3 days co- cultivation) for the two *A. agrestis* isolates under two different MES concentrations in the transformation buffer (initial pH set to 5.5 using the *GV3101 Agrobacterium* strain). *G-I:* Schematic representation of constructs for the expression of two transcription units (TU): one TU for the expression of the *hygromycin B phosphotransferase* (*hph*) gene under the control of the *AaEf1a* promoter and one TU for the expression of G) *p-AaEf1a::hph - p- 35S_s::eGFP-Lti6b* H) *p-AaEf1a::hph - p-AaEF1a::eGFP-Lti6b* and I) *p-AaEf1a::hph - p- AaTip1;1::eGFP-Lti6b*. *J-O:* Confocal images of transformed lines: J) *A. agrestis* Bonn transformed with the *p- AaEf1a::hph - p-AaTip1;1::eGFP-Lti6b* construct. K) *A. agrestis* Bonn transformed with the *p- AaEf1a::hph - p-35S_s::eGFP-Lti6b* construct. L) *A. agrestis* Bonn transformed with the *p- AaEf1a::hph - p-AaEF1a::eGFP-Lti6b* construct. M) *A. agrestis* Oxford transformed with the *p-AaEf1a::hph - p-AaTip1;1::eGFP-Lti6bP* construct. N) *A. agrestis* Oxford transformed with the *p-AaEf1a::hph - p-35S_s::eGFP-Lti6b* construct. O) *A. agrestis* Oxford transformed with the *p-AaEf1a::hph - p-AaEF1a::eGFP-Lti6b* construct. Apical cells of the thallus margin and the surrounding tissue are marked with white arrows. Scale bars: 150 μm. All construct maps at Suppl Table 1. P) Western blot analysis of eGFP accumulation in transgenic lines. Total cellular proteins were separated by polyacrylamide gel electrophoresis, blotted and probed with anti-GFP and anti-actin antibodies. Numbering above blot images corresponds to the identifier of independent transformed lines.

To estimate the transformation efficiency of the protocol for the two *A. agrestis* isolates (Bonn and Oxford), we performed transformation trials using 200 mg of tissue per trial, co-cultivation media with 20mM, 30mM or 40mM MES, *AGL1* or *GV3101 Agrobacterium* harbouring the *p-AaEf1a::hph - p-AaEf1a::eGFP-Lti6b* construct. The number of successful transformants (plant thalli) per experiment is summarised in Fig. 2E and F. We could recover up to three times more transformants for the Oxford isolate compared to the Bonn isolate.

### Characterisation of the activity of the *AaTip1;1* promoter

Previously, a 1368 bp putative promoter fragment of the *Arabidopsis thaliana Tip1;1* gene homolog in *A. agrestis* was selected as a candidate for a constitutive hornwort promoter (Frangedakis, Waller, et al. 2021). Here we further characterise the *AaTip1;1* promoter and compared it with the two other constitutive, *AaEf1a* and the CaMV 35S, promoters used for *A. agrestis*.

Expression of eGFP driven by the *AaTip1;1* promoter is equally strong across the thallus (Fig 2G-O). As reported earlier (Frangedakis, Waller, et al. 2021) expression of eGFP driven by the CaMV 35S promoter seems to be weaker in meristematic areas of the thallus (Fig 2G- O), unlike expression of eGFP driven by the *AaEF1a* promoter which is stronger in meristematic areas of the thallus. It must be noted that the eGFP expression in the stable transformants can be categorised into four groups: Expression throughout the thallus, expression in the rhizoids, expression in patches, or, no expression (Supp Fig 4). Frequency of each of those four events is summarised in Table 1.

We compared the amount of eGFP protein accumulated in plants expressing eGFP under the *AaTip1;1*, the CaMV 35S or the *AaEF1a* promoter. Western blot analysis showed that the *AaTip1;1* is the strongest promoter followed by the *AaEF1a* and the CaMV 35S promoters (Fig 2P).

We also tested the expression of the three constitutive promoters at the sporophyte stage of *A. agrestis* Bonn. When eGFP was driven by the CaMV 35S promoter no eGFP fluorescence could be observed in the sporophyte (Fig 3A, D and G). In contrast, when eGFP expression was driven by the *AaEf1a* or the *AaTip1;1* promoter, eGFP fluorescence was detectable in most sporophyte tissues (Fig 3B, C, E, F, H, I and J).

**Figure 3:**
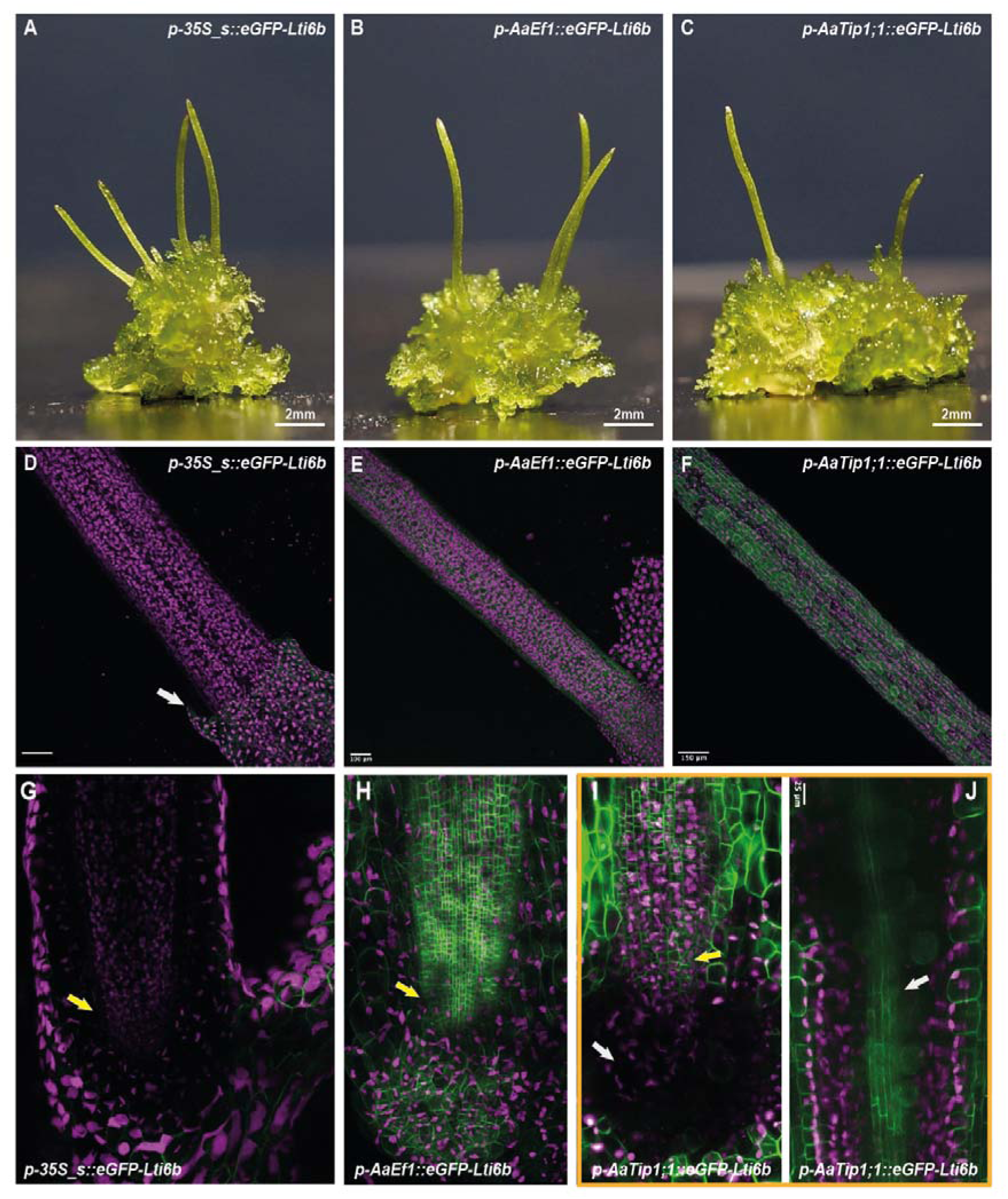
Promoter activity in the *A. agrestis* Bonn sporophyte. A) Transgenic *A. agrestis* Bonn with sporophytes transformed with the *p-AaEf1a::hph - p- 35S_s::eGFP-Lti6b* construct. B) Transgenic *A. agrestis* Bonn with sporophytes transformed with the *p-AaEf1a::hph - p-AaEF1a::eGFP-Lti6b* construct. A) Transgenic *A. agrestis* Bonn with sporophytes transformed with the *p-AaEf1a::hph - p-AaTip1;1::eGFP-Lti6b* construct. *D- F:* Confocal microscopy images (whole-mount) of *A. agrestis* Bonn sporophytes transformed with D) the *p-AaEf1a::hph - p-35S_s::eGFP-Lti6b* construct, Scale bar: 150 μm, and E) the *p-AaEf1a::hph - p-AaEF1a::eGFP-Lti6b* construct. F) the *p-AaEf1a::hph - p- AaTip1;1::eGFP-Lti6b* construct. Scale bar: 150 μm. G) Confocal microscopy images of the sporophyte base of *A. agrestis* Bonn transformed with the *p-AaEf1a::hph - p-35S_s::eGFP-Lti6b* construct, basal meristem indicated with white arrow. Scale bar: 100 μm H) Confocal microscopy images of the sporophyte base of *A. agrestis* Bonn transformed with the *p-AaEf1a::hph - p-AaEF1a::eGFP-Lti6b*, basal meristem indicated with white arrow. Scale bar: 100 μm I- J: Multiphoton microscopy images (whole mount) of the sporophyte base of *A. agrestis* Bonn transformed with the *p-AaEf1a::hph - p-AaTip1;1::eGFP-Lti6b* construct. I) no eGFP signal is visible in the foot (white arrow), but in overlying cell rows of the basal meristem (yellow arrow). J) eGFP signal is strongest in the epidermal cells and the columella (white arrow) Scale bar: 25 μm. All construct maps at Supp Table 1.

### The protocol can be used to transform more hornwort species

Hornworts comprise 11 genera (Villarreal and Renner 2012) which include: *Leiosporoceros, Anthoceros*, *Folioceros*, *Paraphymatoceros*, *Phaeoceros, Notothylas*, *Phymatoceros*, *Phaeomegaceros*, *Nothoceros*, *Megaceros* and *Dendroceros.* (Fig. 4A). We tested whether the optimised protocol for the *A. agresis* isolates can successfully be used to obtain transgenic *A. punctatus* (the model for cyanobacteria symbiosis studies), *L. dussii* (sister to all other hornworts) and *P. carolinianus* plants.

**Figure 4:**
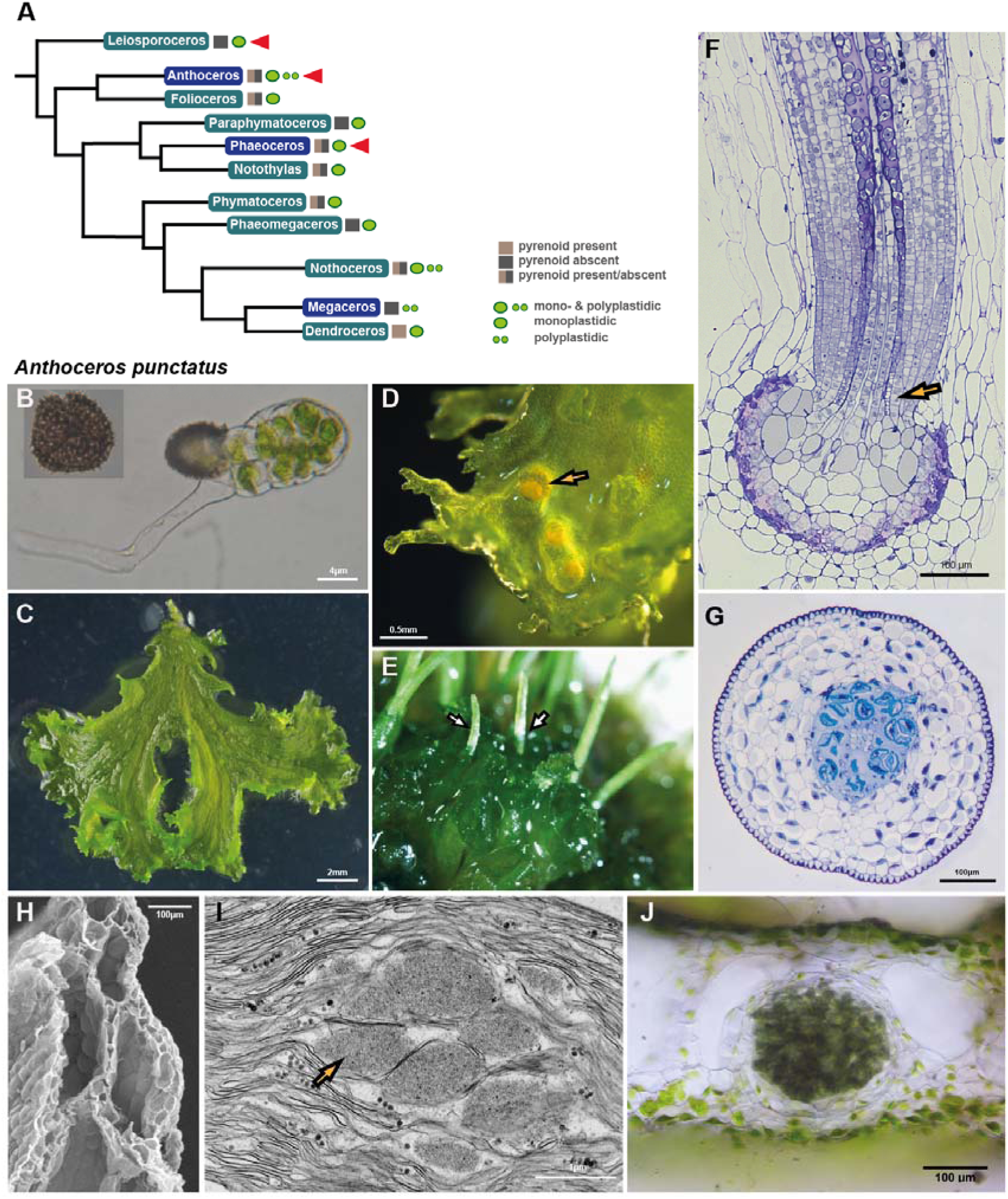
Hornwort phylogeny and the morphological features of *A. punctatus*. A) Hornwort phylogeny (Villarreal and Renner 2012). Genera whose species could be successfully transformed using our improved protocol are marked with red arrowheads. B) Germinating spore which develops into a thallus. Scale bar: 4 μm. C) The thallus is irregularly shaped, lacks specialised internal differentiation and is composed of mucilage chambers and parenchyma cells. Scale bar: 2mm. The male (antheridia, see yellow arrowhead) (D) and female (archegonia) reproductive organs are embedded in the thallus. Scale bar: 0.5 mm. E) Thallus with sporophytes (see white arrowhead). F) Longitudinal section of the foot (see yellow arrowhead) surrounded by the involucre showing the basal meristem and differentiating sporogenous tissue. Scale bar: 100 μm. G) Transverse section of the sporophyte showing its morphology. From the centre to outside: columella, spores, pseudoelaters, assimilative tissue, epidermis and stomata with substomatal cavities. Scale bar: 100 μm. H) SEM of thallus. Scale bar: 100 μm. I) Transmission electron microscopy of the chloroplast (yellow arrowhead: pyrenoid). Scale bar: 1 μm. J) Hand section of *A. punctatus* thallus showing ellipsoidal cavity colonised by cyanobacteria. Scale bar: 100 μm.

*A. punctatus,* as with *A. agrestis,* can routinely be propagated vegetatively in lab conditions, with sub-culturing on a monthly basis. Its genome is sequenced and has a size of approximately 132 Mbp. However, sexual reproduction in lab conditions has been challenging. Spores are punctuated (Fig. 4B). *A. punctatus* thallus is irregularly shaped (Fig. 4C), lacks specialised internal differentiation and is composed of mucilage chambers and parenchyma cells (Fig. 4H). The male (Fig. 4D) and female reproductive organs are embedded in the thallus. Sporophytes grow on the gametophyte (Fig. 4E) and contain columella, spores, pseudoelaters, assimilative tissue, epidermis and stomata (Fig. 4F and G). *A. punctatus* chloroplasts have pyrenoids (Fig. 4I) and its thallus has ellipsoidal cavities colonised by cyanobacteria (Fig. 4J).

*L. dussii* has typically solid thallus with schizogenous cavities in older parts (Fig. 5A). Antheridia are numerous (Fig. 5B and C). Sporophytes (Fig. 5D) develop on the gametophyte. Unlike other hornworts, spores are monolete not trilete and smooth (Fig. 5E). The sporophyte (Fig. 5F) is consisting of the columella, the sporogenous tissue, the assimilative layer, epidermis and stomata. It has a single chloroplast per cell that lacks a pyrenoid (Fig. 5G and H). Chloroplasts have numerous channel thylakoids and extensive grana stacks (Fig. 5I and J). The thallus of *L. dussii* is colonised with Nostoc cyanobacteria (Fig. 5K) which are longitudinally oriented strands in mucilage-filled schizogenous canals.

**Figure 5:**
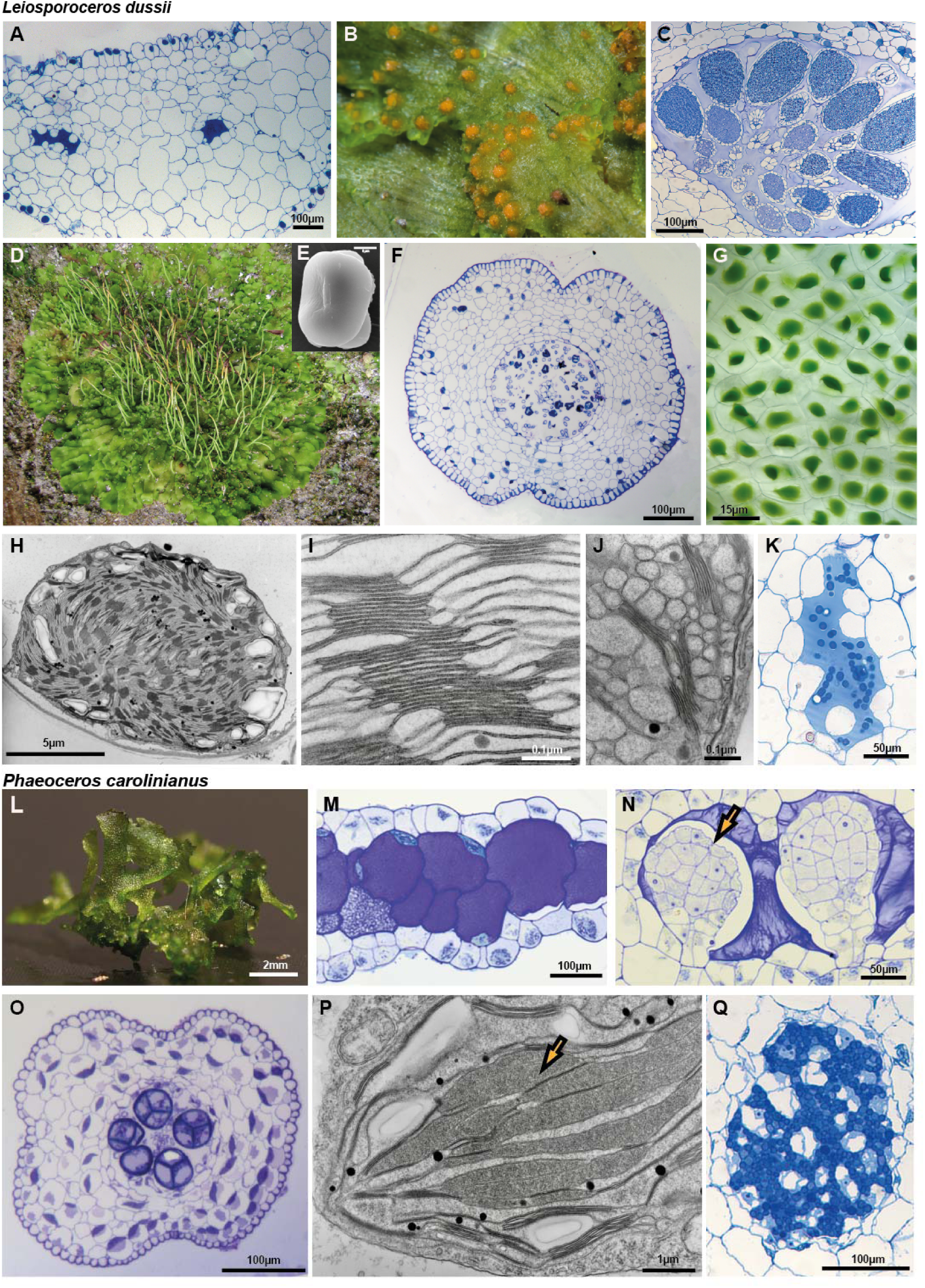
Morphology of Leiosporoceros dussii and Phaeoceros carolinianus. A) *L. dussii* thallus is irregularly shaped, lacks specialised internal differentiation and is composed of mucilage chambers and parenchyma cells. Scale bar: 100 μm. B) *L. dussii* plant with antheridia. C) Light microscopy section showing antheridia embedded in thallus. Scale bar: 100 μm. D) *L. dussii* with sporophytes. E) SEM of spore. Unlike other hornworts it is monolete and smooth. Scale bar: 5 μm. F) Transverse section of the sporophyte showing its morphology. From the centre to outside: columella, spores, pseudoelaters, assimilative tissue, epidermis and stomata with substomatal cavities. Scale bar: 100 μm. G) Light microscopy image of *L. dussii* gametophyte showing a single chloroplast per cell. Scale bar: 15 μm. H-J) Transmission electron microscopy of chloroplast. Scale bar: 5μm (H), 0.1μm (I) and 0.1μm (J). K) Light microscopy section of *Nostoc* colony showing algal cells. Scale bar: 50 μm. L) *P. carolinianus* gametophyte grown under laboratory conditions. Scale bar: 2 mm. M) The thallus with mucilage cells. Scale bar: 100 μm. N) Light microscopy section showing antheridia (yellow arrow) embedded in thallus. Scale bar: 50 μm. O) Transverse section of the sporophyte showing its morphology. From the centre to outside: columella, spores, pseudoelaters, assimilative tissue, epidermis and stomata with substomatal cavities. Scale bar: 100 μm. P) Transmission electron microscopy of chloroplast (yellow arrow pointing to the pyrenoid). Scale bar: 1 μm. Q) Light microscopy section of *Nostoc* colony showing algal cells. Scale bar: 100 μm.

*P. carolinianus* thallus is irregularly shaped (Fig. 5L), lacks specialised internal differentiation (Fig. 5M). It is monoicous and produces both antheridia (Fig. 5N) and archegonia on the dorsal side of the thallus. The sporophyte (Fig. 5O) is consisting of the columella, the sporogenous tissue, the assimilative layer, epidermis and stomata. It has a single chloroplast per cell with pyrenoids (Fig. 5P). The thallus is colonised by cyanobacteria (Fig. 5Q).

We performed preliminary trials using as starting material 200 mg of tissue per trial, co-cultivation media with 30mM or 40mM MES, *AGL1 Agrobacterium* harbouring the *p- AaEf1a::hph - p-AaEf1a::eGFP-Lti6b* plasmid. We successfully recovered stable transformants for all three species (Fig. 6A-F). We also performed trials for the three species using 200 mg of tissue per trial, co-cultivation media with 40mM MES and the *GV3101 Agrobacterium* harbouring the *p-AaEf1a::hph - p-35S_s::eGFP-Lti6b* plasmid, however, we only recovered transformants for *A. punctatus*. The number of successful transformants (plant thalli) per experiment are summarised in Fig. 6G and H. *A. punctatus* has the highest transformation efficiency and *L. dussii* has the lowest efficiency with only three stable lines recovered in total.

**Figure 6:**
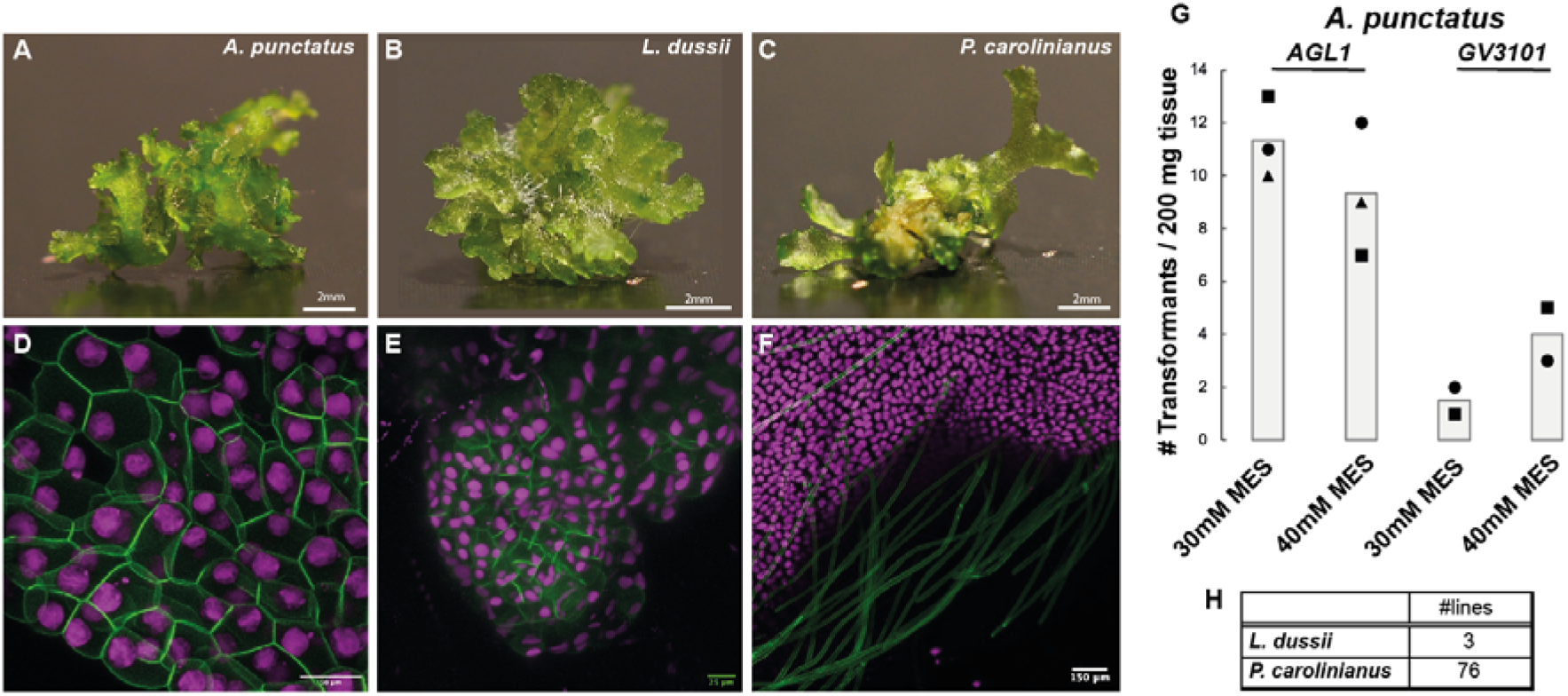
Confocal microscopy images of transgenic *A. punctatus*, *L. dussii* and *P. carolinianus*. *A-C:* Images of *A. punctatus* (A), *L. dussii* (B) and *P. carolinianus* (C) fragments used as starting material for the co-cultivation with *Agrobacterium.* Scale bars: 2 mm. *D-E:* Confocal microscopy image of *A. punctatus* (D) Scale bar: 50 μm, *L. dussii* (E) and *P. carolinianus* (F) gametophytes transformed with the *p-AaEf1a::hph - p-AaTip1;1::eGFP-Lti6b* (D and F) and the *p-AaEf1a::hph - p-AaEf1a::eGFP-Lti6b* construct (E). Scale bar: 150 μm. G) Comparison of number of transformants (per 200 mg starting tissue) (after 3 days co-cultivation) for *A. punctatus* under two different MES concentrations in the transformation buffer (initial pH set to 5.5) using the *AGL1* or *GV3101 Agrobacterium* strain. H) Total number of stable lines obtained for *L. dussii* and *P. carolinianus*.

### The simplified new transformation method

In summary the steps of the new optimised protocols are as follow (Fig. 7, detailed description is provided in Methods and Supp Figures 5, 6, 7 and 8): Fragmented regenerating thallus tissue (grown for at least four weeks) was co-cultivated with *Agrobacterium* in liquid KNOP supplemented with 1% (w/v) sucrose medium. Liquid media were supplemented with 40 mM MES and 3,5-dimethoxy-4-hydroxyacetophenone (acetosyringone) at a final concentration of 100 μM. The pH was adjusted to 5.5. Co-cultivation duration was 3 days with shaking at 21°C on a shaker without any light supplementation (only ambient light from the room). After co-cultivation the tissue was plated on solid KNOP plates supplemented with 100 μg/ml cefotaxime and 10 μg/ml Hygromycin. A month later successful transformants based on rhizoid production were visible on the plate. After 4 weeks it is recommended to transfer the tissue on fresh selective media plates.

**Figure 7:**
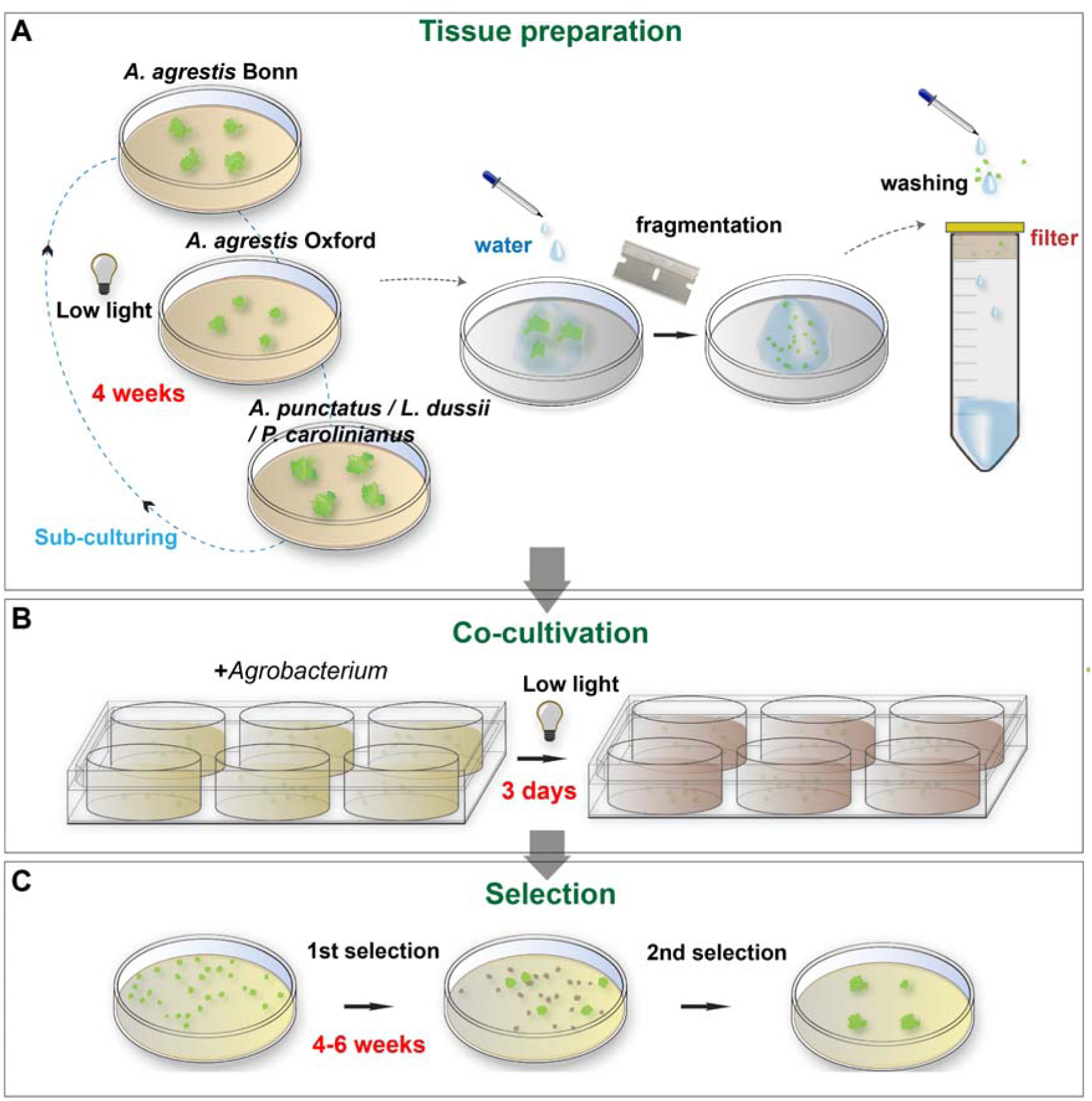
Schematic representation of the optimised transformation protocol. A) Thallus tissue is routinely propagated on a monthly basis under low light. 4-6 week old tissue is fragmented with the aid of a razor blade, transferred to a cell strainer, and washed thoroughly with sterile water. B) The tissue is then co-cultivated with *Agrobacterium* for three days (under low light) and C) spread on antibiotic-containing growth medium. After approximately 4-6 weeks, putative transformants are visible. A final round of selection is used to eliminate false-positive transformants.

### New tools to label and target specific compartments of the hornwort cell

Hornwort cell biology is poorly understood. In order to develop tools to facilitate the study of hornworts at a cellular level, we tested a series of potential signal peptides for subcellular localisation at the mitochondria, Golgi, peroxisome, actin cytoskeleton, chloroplast and the ER (Fig. 8A, Supp Figure 9). We further tested the applicability in hornworts, of new fluorescent proteins, a different reporter and a new selection marker.

**Figure 8:**
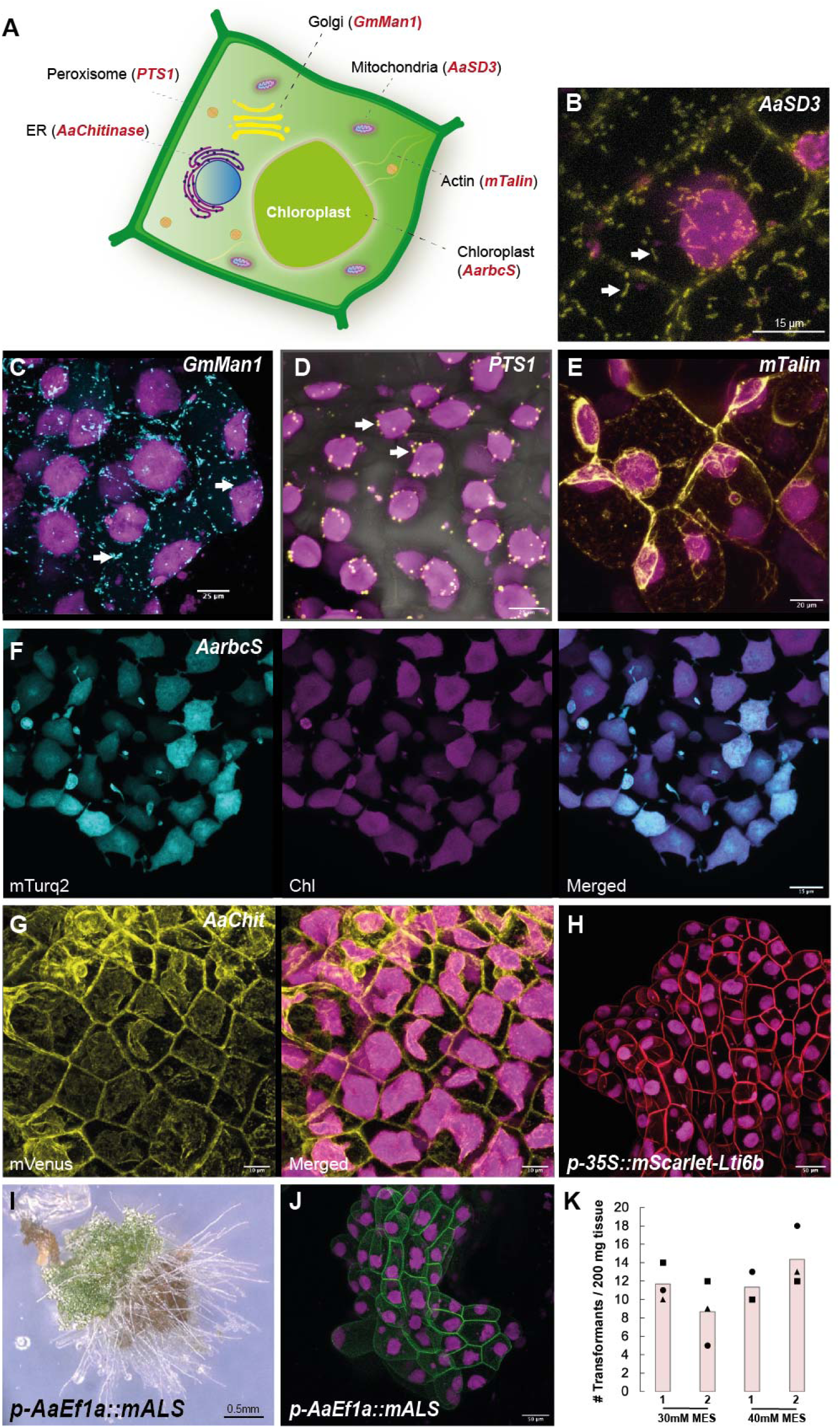
Targeting various subcellular components of the *A. agrestis* thallus with different localization tags. A) Schematic representation of a hypothetical *A. agrestis* cell showing a summary of the new subcellular localisation peptides developed in this study. B) Confocal microscopy image of *A. agrestis* Oxford expressing the *p-AaEf1a::hph - p-AaEF1a::mVenus-AaSD3* construct. Scale bar: 15 μm. C) Confocal microscopy image of *A. agrestis* Oxford expressing the *p- AaEf1a::hph - p-AaEF1a::mTurquoise2-GmMan1* construct. Scale bar: 25 μm. D) Confocal microscopy image of *A. agrestis* Oxford expressing the *p-35S::hph - p-35Sx2::mVenus- PTS1* construct. Scale bar: 25 μm. E) Confocal microscopy image of *A. agrestis* Oxford expressing the *p-35S::hph - p-35S::mVenus-mTalin* construct. Scale bar: 20 μm. F) Confocal microscopy image of *A. agrestis* Oxford expressing the *p-AaEf1a::hph - p-AaEF1a::AarbcS- mTurquoise* construct. Scale bar: 15 μm. G) Confocal microscopy image of *A. agrestis* Oxford expressing the *p-AaEf1a::hph - p-AaEF1a::mVenus-AaChit* construct. Scale bar: 10 μm. H) Confocal microscopy image of *A. agrestis* Oxford expressing the *p-AaEf1a::hph - p- 35S::mScarlet-Lti6b* construct. Scale bar: 50 μm. I) Light microscopy image of 6-week old *A. agrestis* Bonn regenerating fragment transformed with the *p-AaEf1a::mALS - p- 35S_s::eGFP-Lti6b* construct. Scale bar: 0.50 m. J) Confocal microscopy image of *A. agrestis* Bonn expressing the *p-AaEf1a::mALS - p-35S_s::eGFP-Lti6b* construct. Scale bar: 50 μm. K) Comparison of number of transformants (per 200 mg starting tissue) (after 3 days co-cultivation) for *A. agrestis* Bonn under two different MES concentrations in the transformation buffer (initial pH set to 5.5) using the *AGL1* or *GV3101 Agrobacterium* strain harbouring the *p-AaEf1a::mALS - p-35S_s::eGFP-Lti6b* construct.

Mitochondria in the liverwort *Marchantia polymorpha* have been visualised using a targeting sequence derived from the *A. thaliana Segregation Distortion 3* (*SD3)* gene, which encodes a protein with high similarity to the yeast translocase of the inner mitochondrial membrane 21 (TIM21) (Ogasawara et al. 2013). A mitochondrial targeting sequence (MTS) from the *Saccharomyces cerevisiae COX4* (*ScCOX4*) gene (Hamasaki et al. 2012) can also be used successfully in *M. polymorpha* (Supp Figure 10). However, mitochondrial localisation was not achieved when using either the *A. thaliana SD3* or the *ScCOX4* targeting sequences in *A. agrestis* Oxford. We thus tested the N-terminal sequence of a predicted *SD3 A. agrestis* gene (Sc2ySwM_117.1379.1) (Supp Figure 9), for its ability to direct fluorescent localisation in mitochondria. Using this sequence (*p-AaEf1a::hph - AaEF1a::mVenus-AaSD3* construct), we observed fluorescence in structures in the cytosol that resemble mitochondria (Fig. 8B). However, it must be noted that the signal has low fluorescence intensity and the use of a confocal microscope at high magnification is necessary for observation.

In *M. polymorpha* the Golgi has been visualised using the transmembrane domain of the rat sialyltransferase (ST) as a targeting sequence (Kanazawa et al. 2016). Another Golgi localisation sequence which can be used in *M. polymorpha* is a targeting sequence derived from the soybean *(Glycine max)* α*-1,2 mannosidase 1* (*GmMan1)* gene (Luo and Nakata 2012) (Supp Figure 9). We tested whether the ST or the *GmMan1* peptides can be successfully used in *A. agrestis* Oxford for Golgi targeting. Only the *GmMan1* peptide (*p- AaEf1a::hph - p-35Sx2::mTurquoise2-GmMan1* construct) led to fluorescence in structures in the cytosol that resemble Golgi (Fig. 8C).

Previous studies in *M. polymorpha* have used the peroxisomal targeting signal 1 (PTS1) (Ser-Lys-Leu) as a peroxisome targeting sequence (Ogasawara et al. 2013) (Supp figure 9). We tested whether the PTS1 signal peptide (*p-35S::hph - p-35Sx2::mVenus-PTS1* construct) can be successfully used in *A. agrestis* Oxford for peroxisome targeting. We observed fluorescence in structures in the cytosol that resemble peroxisomes (Fig. 8D).

The C-terminus of mouse talin (mTalin) has been used for actin labelling in *Arabidopsis* and *M. polymorpha* (Kost, Spielhofer, and Chua 1998; Kimura and Kodama 2016). We generated *A. agrestis* Oxford plants stably transformed with a *p-35S::hph - p35S::mVenus-mTalin* construct and we observed fluorescence localisation in the filamentous structures in the cytosol (Fig. 8E).

The chloroplast transit peptide of the Rubisco small subunit (rbcS) has been used for protein targeting to the chloroplast in angiosperms (Kim et al. 2010; Shen et al. 2017). We cloned the predicted chloroplast transit peptide of the *A. agrestis* rbcS (Sc2ySwM_344.2836.1) (Supp Figure 9). When fused with mTurquoise2 (*p-AaEf1a::hph - AaEF1a::AarbcS- mTurquoise2* construct) we observed fluorescence localisation in the chloroplasts (Fig. 8F).

In *M. polymorpha* the N-terminal targeting sequence from a predicted chitinase (Mp2g24440) in combination with the C-terminal HDEL ER retention peptide, has been successfully used for ER localisation (Sauret-Güeto et al. 2020). We tested the N-terminal targeting sequence of a predicted chitinase in *A. agrestis* (Sc2ySwM_228.5627.1) (Supp Figure 9), for its ability to direct fluorescent localisation in ER (*p-AaEf1a::hph - AaEF1a::mVenus-AaChit* construct). We observed fluorescence in reticulate structures in the cytosol around the nuclei and in the plasma membrane (Fig. 8G).

Finally, we tested the targeting sequence derived from the *A. thaliana Calcineurin B-like 3* (*AtCBL3)* gene for Tonoplast localisation, that is functional in *M. polymorpha* (Supp Fig 10). However, we were not able to detect any fluorescent signal in *A. agrestis* Oxford.

We also confirmed that the tags targeting mitochondria, Golgi, chloroplast, and ER can be also used for *A. punctatus* however the signal is not as strong as in *A. agrestis* (Supp Figure 11). Summary of lines obtained for both *A. agrestis* and *A. punctatus* in Table 2.

**Table 2:** Summary of transgenic lines expressing different organelle targeting constructs.

|  | *construct* | *Species/strain* | *Number of transformants* | *Expression in rhizoids* | *Patchy expression* | *No fluorescence* |
| --- | --- | --- | --- | --- | --- | --- |
|  | *p-AaEf1a::hph - AaEF1a::mVenus-AaSD3* | *Oxford* | 5 | - | 1 | 1 |
|  | *p-AaEf1a::hph - p-35Sx2::mTurquoise2-GmMan1* | *Oxford* | 15 | 2 |  | 2 |
|  | *p-35S::hph - p-35Sx2::mVenus-PTS1* | *Oxford* | 17 | 2 | 4 | 1 |
|  | *p-35S::hph - p-35S::mVenus-mTalin* | *Oxford* | 5 | - | - | - |
|  | *p-AaEf1a::hph - AaEF1a::AarbcS-mTurquoise2* | *Oxford* | 14 | 2 | - | - |
|  | *p-AaEf1a::hph - AaEF1a::mVenus-AaChit* | *Oxford* | 5 | - | - | 1 |
|  | *p-AaEf1a::hph - p-35Sx2::mVenus-GLS-ST* | *Bonn* | 5 | - | - | 5 |
|  | *p-AaEf1a::hph - AaEF1a::mVenus-AaSD3* | *A. punctatus* | 5 | - | - | 1 |
|  | *p-AaEf1a::hph* - *p-35Sx2::mTurquoise2-GmMan1* | *A. punctatus* | 5 | - | - | 1 |
|  | *p-AaEf1a::hph* - *AaEF1a::AarbcS-mTurquoise2* | *A. punctatus* | 6 | - | - | 1 |
|  | *p-AaEf1a::hph* - *AaEF1a::mVenus-AaChit* | *A. punctatus* | 5 | - | - | 1 |
|  | *p-AaEF1a::hph* - *p-35S::mScarlet-Lti6b* | *Bonn* | 31 | 1 | 1 | 9 |

#### Applicability of the fluorescent protein mScarlet in hornworts

To expand the palette of fluorescent proteins (FP) that can be used in *A. agrestis*, we tested the expression of the synthetic monomeric Scarlet (mScarlet) “red” fluorescent (Bindels et al. 2017). mScarlet is the brightest among the monomeric red FP with a greater than 3.5 times brightness increase compared to mCherry for example. Given that its emission maxima is 594 nm it can be combined with other FP such as eGFP, mVenus or mTurquoise2 with minimal spectral overlap. We successfully generated *A. agrestis* Bonn lines expressing mScarlet (Fig. 8H). Summary of lines obtained in Table 2.

#### Applicability of the Ruby reporter and 2A peptides in hornworts

RUBY is a new reporter (He et al. 2020) that converts tyrosine to red colour betalain that is visible without the need for extra processing steps. Three betalain biosynthetic genes (*CYP76AD1*, *DODA* and *Glucoysl transferase*) are fused into a single transcription unit using a single promoter the P2A self-cleavage peptides and a terminator. We generated a construct where we fused the *AaTip1;1* promoter with the RUBY cassette and the double NOS-35S terminator. However, when the plants transformed with the construct were examined, we did not observe any red colour (Supp Figure 12A).

2A self-peptides are known to have different cleavage efficiencies (Liu et al. 2017). To test the efficiency of 2A self-cleavage peptides we generated two constructs where the mVenus FP fused to a nucleus targeting sequence and the mTurquoise2 FP fused to a plasma membrane targeting sequence were combined into a single transcription unit separated by either the P2A peptide or the E2A (Supp figure 12B-I). When used in *M. polymorpha* the expected FP localisation was observed (Supp figure 12J and K). However, in the case of *A. agrestis* Oxford mainly mVenus localisation into the nucleus was observed suggesting that cleavage efficiency is very low. Summary of lines obtained in Supp Table 2.

### New selection marker

The acetolactate synthase (ALS) is one of the enzymes that catalyse the biosynthesis of three essential amino acids, leucine, isoleucine and valine. The herbicide chlorsulfuron is an ALS inhibitor and has been used in combination with mutated *ALS* genes as a selection marker in different plant species (Kawai et al. 2008). We first tested whether *A. agrestis* is susceptible to chlorsulfuron and found that a 3 weeks incubation period with 0.5 μM chlorsulfuron was sufficient to inhibit growth of untransformed thallus tissue (Supp Figure 13). We then performed trials where we used a *M*. *polymorpha* mutated *ALS* gene (Ishizaki et al. 2015) driven by the *AaEf1a* promoter as a selectable marker, the *GV3101 Agrobacterium* strain and 30 or 40mM MES. Up to 18 chlorsulfuron resistant plants were successfully recovered from 200 mg of tissue (Figure 8I, J and K). It must be noted that growth of transgenic plants on chlorsulfuron selection media is slower compared to hygromycin. It takes approximately an additional two weeks until primary transformants are visible by naked eye.

### Conclusions

After testing several different approaches to increase transformation efficiency in the hornwort *A. agrestis*, it was pH control during co-cultivation that proved most successful. Our results suggest that a pH of the co-cultivation medium around 5.5 is a key parameter to increase *Agrobacterium* infection rates, similar to what has been reported for *A. thaliana Agrobacterium*-mediated transformation (Wang et al., 2018). Consequently, our results support the notion of conserved innate immune responses via calcium signalling in land plants. These results may help to establish *Agrobacterium*-mediated transformation methods for plants that currently lack an efficient transformation method (e.g. lycophytes or various streptophyte algal lineages).

## Methods

### Plant material and maintenance

In this study we used the *Anthoceros agrestis* Oxford and Bonn isolates (Szövényi et al. 2015) and *Anthoceros punctatus* (Li et al. 2014). Mature gametophytes and sporophytes of *A. punctatus* were originally collected from a glasshouse at the Royal Botanic Garden of Edinburgh, by Dr David Long. *Leiosporoceros dussii* was collected at Río Indio in Panama (N08°38.521’, W80°.06.825, Elev. 801 m) by Juan Carlos Villarreal. *Phaeoceros carolinianus* collected in Louisiana, USA (provided by Fay-Wei Li) and grown under similar axenic conditions with *A. agrestis*.

*A. agrestis, A. punctatus, L. dussii and P. carolinianus* thallus tissue was propagated on KNOP medium (0.25 g/L KH_2_PO_4_, 0.25 g/L KCl, 0.25 g/L MgSO_4_•7H_2_O, 1 g/L Ca(NO_3_)_2_•4H_2_O and 12.5 mg/L FeSO_4_•7H_2_O). The medium was adjusted to pH 5.8 with KOH and solidified using 7.5 g/L Gelzan CM (#G1910, SIGMA) in 92x16 mm petri dishes (#82.1473.001, SARSTEDT) with 25-30 ml of media per plate. Plants were routinely grown in a tissue culture room (21°C, 12 h of light and 12 h of dark, 3-5 or 35 μmol m^−2^ s^−1^ light intensity, Philips TL-D 58W (835)). To subculture the thallus tissue, a small part of it (approximately 2mm x 2mm) was cut using sterile disposable scalpels (#0501, Swann Morton) and placed on fresh media on a monthly basis (Supp Figure 2).

*Marchantia polymorpha* accessions Cam-1 (male) and Cam-2 (female) were used in this study (Delmans, Pollak, and Haseloff 2017). Plants were grown on half strength Gamborg B5 medium plus vitamins (Duchefa Biochemie G0210, pH 5.8) and 1.2% (w/v) agar (Melford capsules, A20021), under continuous light at 22 °C with light intensity of 100 μmol m^−2^ s^−1^.

### Tissue preparation for transformation

For the preparation of tissue used for transformation (fragmentation approach using razor blades), small pieces of thallus, approximately 2mm x 2mm, were cut using sterile disposable scalpels (#0501, Swann Morton) and placed on plates containing fresh growth medium (15-20 fragments per plate). The plates were grown for 4-6 weeks, at 21°C, 12 h of light and 12 h of dark, 3-5 μmol m^−2^ s^−1^ light intensity (Supp Figure 2).

After 4-6 weeks, 1 g of thallus tissue (approximately 10 petri dishes) was transferred into an empty petri dish, 2-10 ml of water were added and then fragmented using a razor blade (#11904325, Fisher Scientific) for approximately 5 mins (Supp Video 1). The fragmented tissue was washed with 100 ml of sterile water using a 100 µm cell strainer (352360; Corning) or until the flow through was clear (see Supp Figure 5, 6, and 8).

When tissue was fragmented using a homogeniser, approximately 1 g of thallus tissue was homogenised in 20 ml sterile water using an Ultra-Turrax T25 S7 Homogenizer (727407; IKA) and corresponding dispensing tools (10442743; IKA Dispersing Element), for 5 sec, using the lowest speed of 8000 rpm. The homogenised tissue was washed with 100 ml of sterile water using a 100 µm cell strainer (352360; Corning) or until the flow through was clear

### *Agrobacterium* culture preparation

One to three *Agrobacterium* colonies (*AGL1* strain) were inoculated in 5 ml of LB medium supplemented with rifampicin 15 μg/ml (#R0146, Duchefa), carbenicillin 50 μg/ml (#C0109, MELFORD) and the plasmid-specific selection antibiotic spectinomycin 100 μg/ml (#SB0901, Bio Basic). For the *GV3101* strain preparation, one to three colonies were inoculated in 5 ml of LB medium supplemented with rifampicin 50 μg/ml (#R0146, Duchefa), gentamicin 25 μg/ml (#G0124, Duchefa) and spectinomycin 100 μg/ml (#SB0901, Bio Basic). The pre-culture was incubated at 28°C for 2 days with shaking at 120 rpm.

### Co-cultivation

Co-cultivation medium was liquid KNOP with 1% (w/v) sucrose (0.25 g/L KH_2_PO_4_, 0.25 g/L KCl, 0.25 g/L MgSO_4_•7H_2_O, 1 g/L Ca(NO_3_)_2_•4H_2_O, 12.5 mg/L FeSO_4_•7H_2_O and 10 g/L sucrose), supplemented with 40 mM MES, with pH 5.5 adjusted with KOH. The medium was filter sterilised (#430767, Corning Disposable Vacuum Filter/Storage Systems) and was stored in 50 ml falcon tubes in -20°C if necessary.

5 ml of a 2 day *Agrobacterium* culture was centrifuged for 7 min at 2000 *g*. The supernatant was discarded, and the pellet was resuspended in 5 ml liquid KNOP supplemented with 1% (w/v) sucrose, (S/8600/60; ThermoFisher, Loughborough, UK), 40 mM MES (255262A, ChemCruz), pH 5.5 and 100 μM 3′,5′-dimethoxy-4′-hydroxyacetophenone (acetosyringone) (115540050; Acros Organics, dissolved in dimethyl sulfoxide (DMSO) (D8418; Sigma). The culture was incubated with shaking (120 rpm) at 28°C for 3-5 h. The fragmented thallus tissue was transferred into the wells of a six-well plate (140675; ThermoFisher) with 5 ml of liquid KNOP medium supplemented with 1% (w/v) sucrose and 40 mM MES and the pH was adjusted to 5.5. Then, 80 μl of *Agrobacterium* culture and acetosyringone at final concentration of 100 μM were added to the medium.

The tissue and *Agrobacterium* were co-cultivated for 3 days with an orbital shaker at 110 rpm at 22°C without any additional supplementary light (only ambient light from the room, 1–3 μmol m^−2^ s^−1^). After 3 days, the tissue was drained using a 100 µm cell strainer (352360; Corning) and moved onto solid KNOP plates (two Petri dishes from a single well) supplemented with 100 μg/ml cefotaxime (BIC0111; Apollo Scientific, Bredbury, UK) and 10 μg/ml hygromycin (10687010; Invitrogen) or 0.5 μM Chlorsulfuron (Sigma 64902-72-3 - to prepare 1000x stock Chlorsulfuron solution, 100mM chlorsulfuron super-stock solution was prepared in DMSO first, and then a 500µM stock solution in dH_2_O).

If necessary, after 4 weeks, plants were transferred to fresh solid KNOP plates supplemented with 100 μg/ ml cefotaxime and 10 μg/ ml hygromycin and grown at 22°C under 12 h light : 12 h dark at a light intensity of 35 μmol m^−2^ s^−1^ (TL-D58W/835; Philips).

### Optimisation trials

To test the effect of MES concentration on the transformation efficiency 3 g of *A. agrestis Bonn* thallus tissue was harvested four weeks after subculturing, fragmented with a razor blade, washed with 100 ml water and distributed into six equal parts of 500 mg. Each part was then placed into a well of a 6-well plate with 5 ml co-cultivation medium with different MES concentrations. The MES concentration of the co-cultivation medium in each of the six wells was: 0, 10, 20, 30, 40 or 50 mM. 100 ul of the same *Agrobacterium* pre-culture, containing a *p-AaEf1a*::*hph* - *p-AaTip1;1-eGFP-Lti6b* construct, was added to each well. Three replicates of this trial were performed.

To test the effect of co-cultivation medium pH value on the transformation efficiency, 2.5 g of *A. agrestis Bonn* thallus tissue was harvested four weeks after subculturing, fragmented with a razor blade, washed with 100 ml sterile distilled water and distributed into 5 equal parts of 500 mg. Each part was then placed into a well of a 6-well plate with 5 ml co-cultivation medium containing 40 mM MES, adjusted to different pH values. The pH value of the co-cultivation medium in each of the five wells was adjusted with 1 M KOH to: 5, 5.25, 5.5, 5.75 or 6. 100 ul of the same *Agrobacterium* pre-culture, containing a *p-AaEf1a::hph - p- AaTip1;1-eGFP-Lti6b* construct, was added to each well. Three replicates of this trial were performed.

To test the effect of co-cultivation sucrose concentration on the transformation efficiency, approximately 1.2 g of *A. agrestis Bonn* thallus tissue was harvested four weeks after subculturing, fragmented with a razor blade, washed with 100 ml water, distributed into six equal parts of 200 mg, and each part was then placed into a well of a 6-well plate. In each plate, the sucrose concentration in three wells was 1% (w/v) and in the remaining three was 2% (w/v). 80 ul of the same *Agrobacterium* (transformed with the *p-AaEf1a::hph - - 35S_s::eGFP-Lti6b* construct) pre-culture was added to each well The experimental set up was similar for testing the effect of the *Agrobacterium* strain (*AGL1* or *GV3101*) on the transformation efficiency as well as the *A. agrestis* Oxford optimisation trials with the only difference that the *Agrobacterium* was transformed with the *p-AaEf1a::hph - p- AaEf1a::eGFP-Lti6b* construct.

Co-cultivation and selection were carried out as described in the previous section. After 3-4 weeks on a first round of selection, surviving and growing thallus fragments were transferred to fresh selection plates. After an additional 2 months on 2nd selection, growing thallus pieces that showed GFP expression (observed with a Leica M205 FA Stereomicroscope, GFP longpass filter) were counted as stable transformants.

### Marchantia polymorpha transformation

Transgenic *M. polymorpha* plants were obtained according to (Sauret-Güeto et al. 2020).

### Construct generation

Constructs were generated using the OpenPlant toolkit (Sauret-Güeto et al. 2020). OpenPlant L0 parts used: OP-019 CDS_mALS, OP-023 CDS12-eGFP, OP-020 CDS_hph, OP-027 CDS12_mTurquoise2, OP-029 CDS12_mVenus, OP-037 CTAG_Lti6b, OP-054 3TER, _Nos-35S and OP-049 PROM_35S (for detailed maps see Supp Table 1). The DNA sequence of target peptides described in this study were synthesised using the Genewiz or the IDT company and cloned into the pUAP4 vector. The information about the sequences of the target peptides used in this study can be found in Supp Figure 9.

### Western blotting

50 mg of *A. agrestis* tissue, grown for 3 weeks on KNOP medium at 21 °C in 12 hours light (5 μmol m^−2^ s^−1^) : 12 hours dark, were placed in 1.5 ml tube with two metal beads, flash frozen in liquid nitrogen and grounded using a TissueLyser II (#85300, Qiagen) at 30Hz for 5 mins. The tissue powder was resuspended in 400 μL 10× Laemmli loading buffer (0.5 M Tris-HCl pH 6.8, 20 % w/v SDS, 30 % v/v glycerol, 1 M DTT, 0.05 % w/v bromophenol blue,) supplemented with Roche cOmplete protease inhibitor (# 11836170001, Roche)), heated at 90 C° for 10 minutes and centrifuged at 10,000 g for 5 minutes. The supernatant was transferred to a new tube. Equal amounts of proteins were separated by denaturing electrophoresis in NuPAGE gel (#NP0322BOX, Invitrogen) and electro-transferred to nitrocellulose membranes using the iBlot2 Dry Blotting System (ThermoFisher). eGFP was immuno detected with anti-GFP antibody (1:4000 dilution) (JL-8, #632380, Takara) and anti-mouse-HRP (1:15000 dilution) (#A9044, Sigma) antibodies. Actin was immuno detected with anti-actin (plant) (1:1500 dilution) (#A0480, Sigma) and (1:15000 dilution) anti-mouse-HRP (1:15000 dilution) (#A9044, Sigma) antibodies, using the iBind™ Western Starter Kit (#SLF1000S, ThermoFisher). Western blots were visualised using the ECL™ Select Western Blotting Detection Reagent (#GERPN2235, GE) following the manufacturer’s instructions. Images were acquired using a Syngene Gel Documentation system G:BOX F3.

### Sample preparation for Imaging

A gene frame (#AB0576, ThermoFisher) was positioned on a glass slide. A thallus fragment was placed within the medium-filled gene frame together with 30 μL of milliQ water. The frame was then sealed with a cover slip. Plants were imaged immediately using a Leica SP8X spectral fluorescent confocal microscope.

### Imaging with Confocal Microscopy

Images were acquired on a Leica SP8X spectral confocal microscope. Imaging was conducted using either a 10× air objective (HC PL APO 10×/0.40 CS2) or a 20× air objective (HC PL APO 20×/0.75 CS2). Excitation laser wavelength and captured emitted fluorescence wavelength window were as follows: for mTurquoise2 (442 nm, 460−485 nm), for eGFP (488 nm, 498−516 nm), for mVenus (515 nm, 522−540 nm), and for chlorophyll autofluorescence (488 or 515, 670−700 nm).

Sporophytes of promoter-eGFP reporter lines were dissected from the gametophyte, and the gametophyte tissue around the base carefully removed with scalpels under a dissecting scope without disturbing the sporophyte base. The sporophytes were then transferred into a Lab-Tek chambered coverglass (#155361, Lab-Tek) and overlaid with 5mm solid KNOP medium (solidified with Gelrite) and water to keep them in place and prevent dessication. eGFP-expression within the sporophyte base of the living sporophytes was then visualised with a Leica TCS SP8 MP, excitation with a multiphoton laser at 976 nm, eGFP fluorescence detected with a bandpass filter (HyD-RLD 2ch FITC filter 525/50) and chloroplast autofluorescence with HyD-RLD 675/55.

### Light microscopy

Images were captured using a KEYENCE VHX-S550E microscope (VHX-J20T lens) or a Leica M205 FA Stereomicroscope (with GFP longpass (LP) filter).

## Acknowledgements

We are thankful to Juan Carlos Villarreal, Laval University for providing *L. dussii cultures* and Fay-Wei Li, Cornell University for providing *P. carolinianus cultures.* Fig. 4C picture credit: John Baker, Oxford University. Fig. 4E: picture credit: Keiko Sakakibara, Rikkyo University. Fig. 4F, K and L: picture credit: Masaki Shimamura, Hiroshima University. Fig. 5B and D: picture credit: Juan Carlos Villarreal.

## Author contributions

MW, EF, and PS conceived and designed the experiments. EF and MW performed optimisations, generated and characterized the hornwort transgenic lines. EF, MW, KZ and AOM performed imaging. JR and SSG generated and characterized the transgenic *M. polymorpha* lines. JH and JMH provided resources. CRRS performed *A. punctatus* optimisations. MW, EF and PS wrote the article with contributions of all the authors.

