## Supplementary material for "An optimised transformation protocol for *Anthoceros agrestis* and three more hornwort species": Supp info

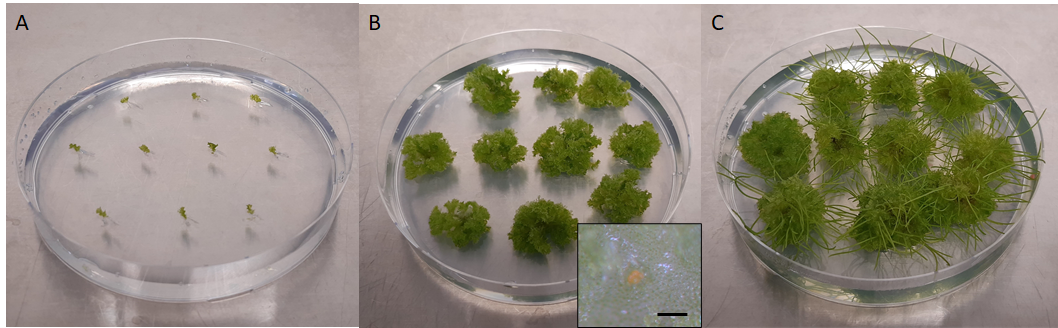

**Figure S1: Sporophyte induction for *A. agrestis* Bonn**

To induce development of sporophytes, *A. agrestis* Bonn thallus fragments were sub-cultured as described in Methods, under Plant material and maintenance (A). The established plant cultures were then kept in a growth chamber at 24h light, 20-30 μmol m^−2^ s^−1^, 60% humidity, 23°C and grown for approx. 1.5 - 2 months, until the individual thallus pieces would grow into clumps of tissue with a diameter of approximately 1-2 cm (B). At this point, the cultures were examined under the stereomicroscope and if antheridia could be observed (B), approx. 5 ml of sterile water was poured over the tissue and added to the plate. The plates were then incubated for another 2-3 weeks in the growth chamber, after which the appearance and growth of sporophytes could be observed (C) (scale bar = 100 μm)

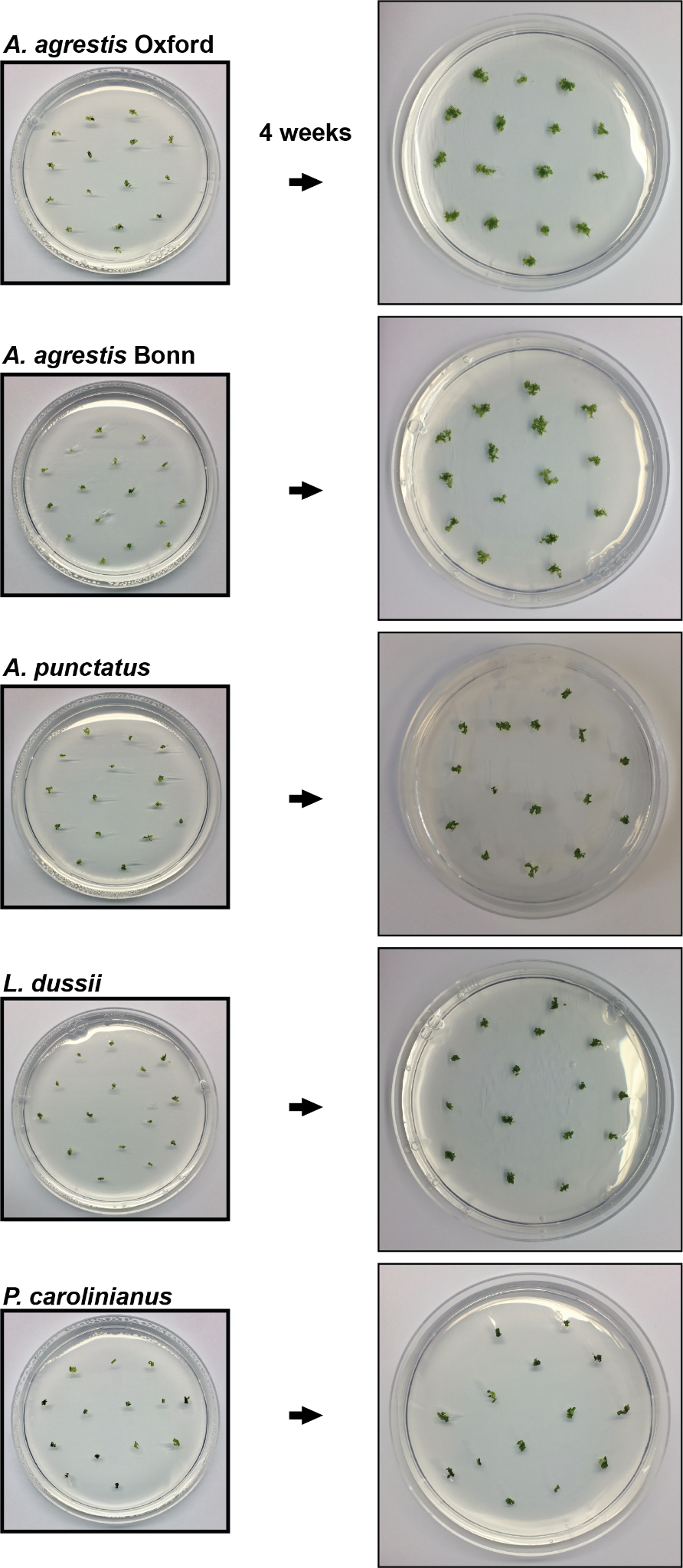

**Figure S2: Tissue culturing for *A. agrestis, A. punctatus, L. dussii and P. carolinianus.***

Images to demonstrate morphology of thallus used for routine tissue propagation for *A. agrestis, A. punctatus, L. dussii and P. carolinianus.*

Petri dish dimensions: 92 x16 mm.

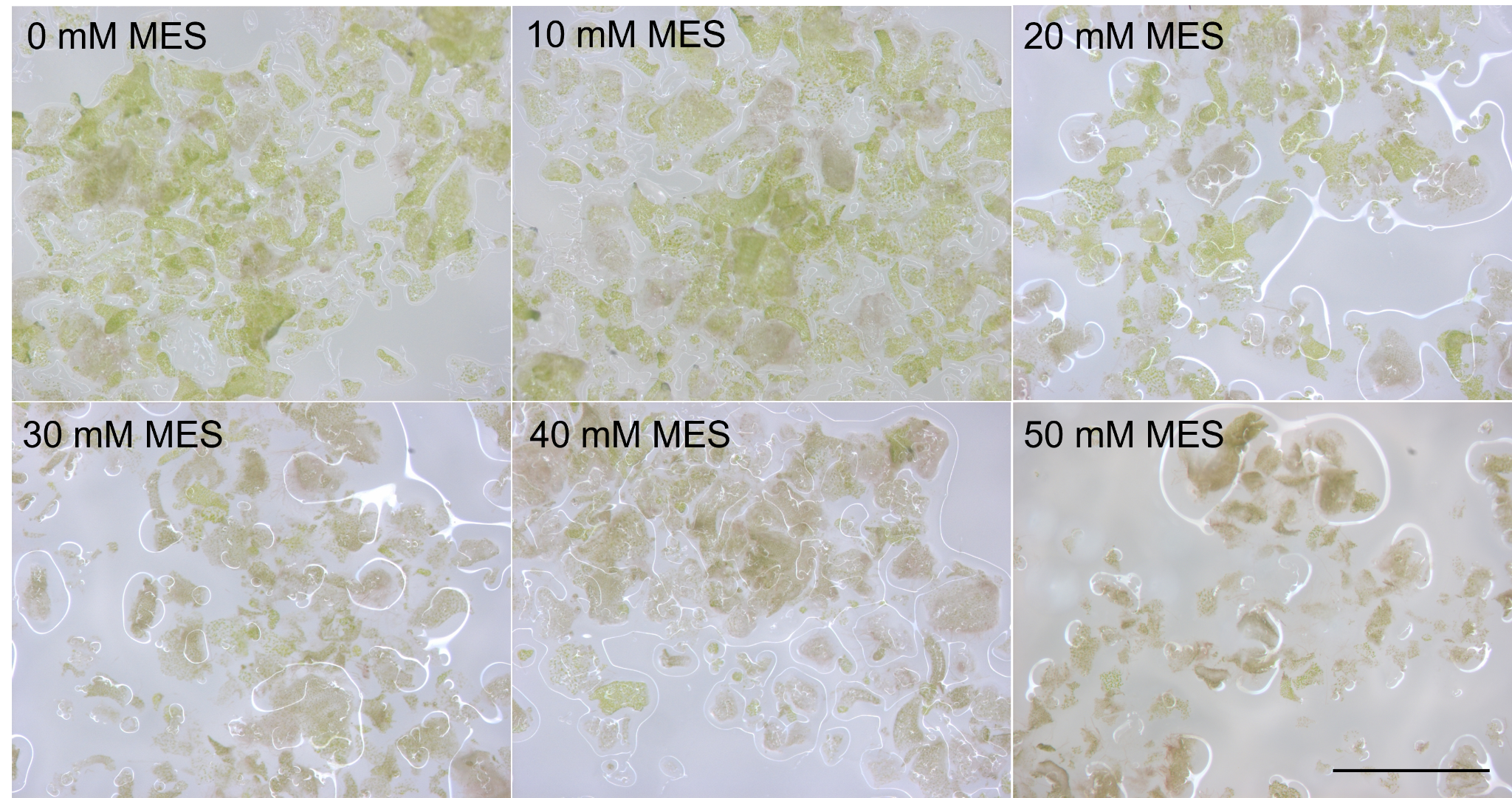

**Figure S3: Effect of different MES concentrations in the transformation buffer on *A. agrestis Bonn* tissue fragments after 3 days of co-cultivation.**

With increased MES concentration, tissue fragments look unhealthy or dead 3 days after co-cultivation as indicated by the brown/grey colour of the chloroplasts. At 50 mM MES, almost no living cells (with green chloroplasts) could be found (scale bar = 1 mm).

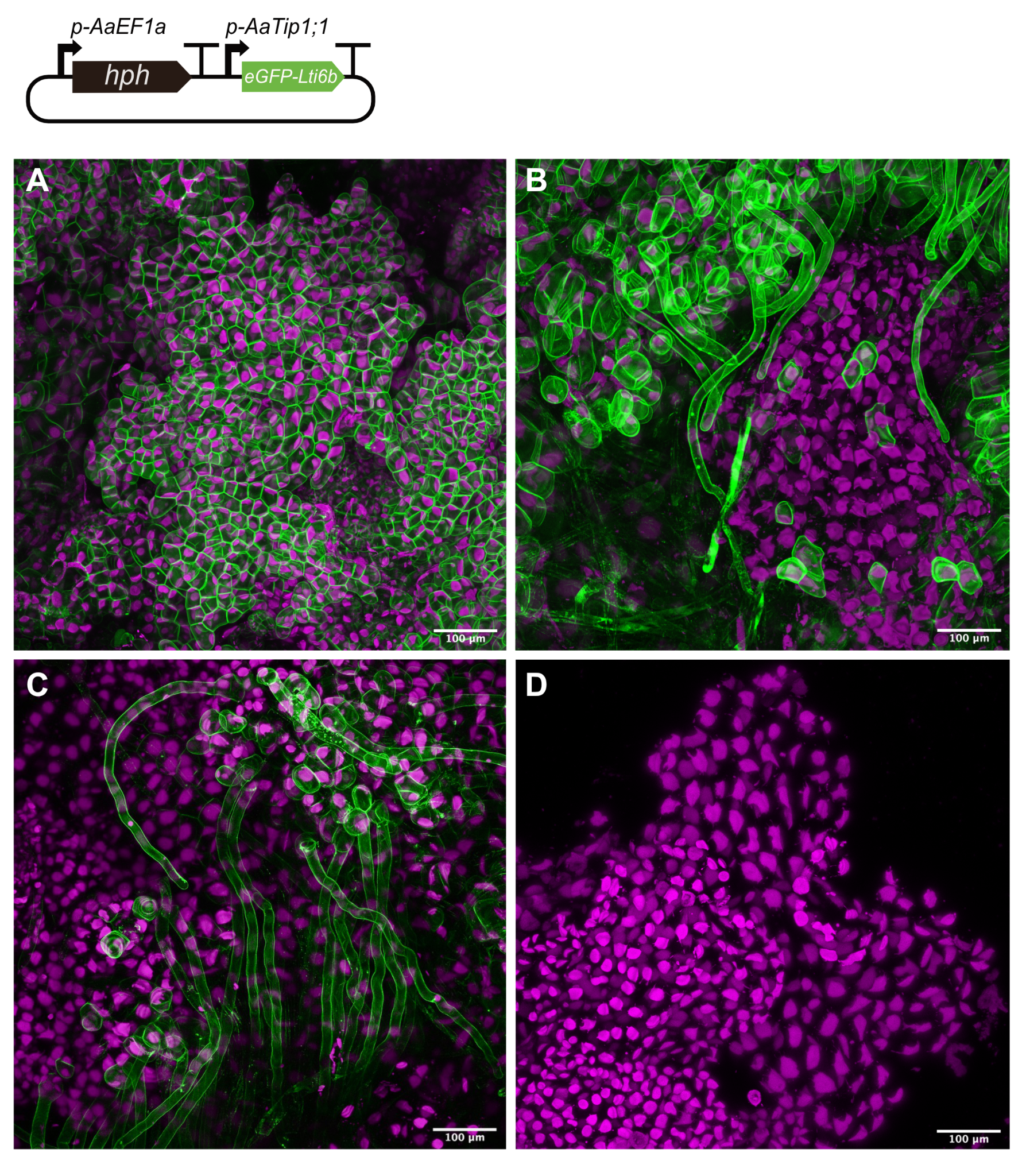

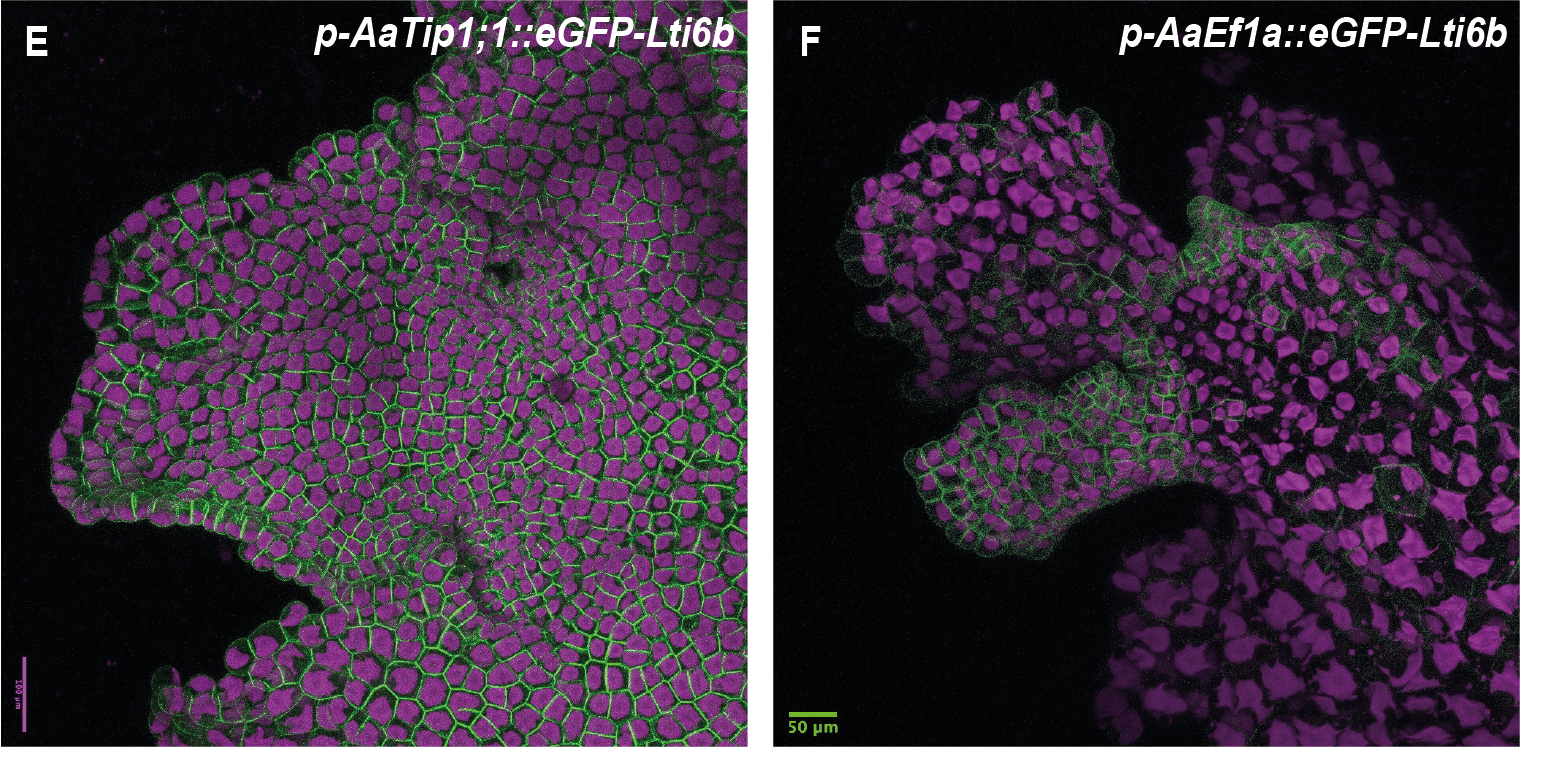

**Figure S4: Activity of the *AaTip1;1* and the *AaE1a* promoters in the gametophyte of *A. agrestis*.**

On the top of the figure: Schematic representation of constructs for the expression of two transcription units (TU): one TU for the expression of the hygromycin B *phosphotransferase* (*hph*) gene under the control of the *AaEf1a* promoter and one TU for the expression of *p-AaTip1;1::eGFP-Lti6b.* A-D) Confocal images of *A. agrestis* Bonn transformed with the *p-AaTip1;1::eGFP-Lti6b* construct. A) Thallus margin. B) Rhizoids and irregular thallus tissue expression. C) Rhizoid only expression. D) Section of the thallus not expressing eGFP. Scale bars: 100 μm. E) Gametophyte thallus margin of *A. punctatus* transformed with the *p-AaEf1a::hph - p-AaTip1;1::eGFP-Lti6b* construct. Scale bar: 100 μm. F) Gametophyte thallus margin of *A. punctatus* transformed with the *p-AaEf1a::hph - p*-*AaE1a::eGFP-Lti6b* construct. Scale bar: 50 μm. All construct maps at Supp Table 1.

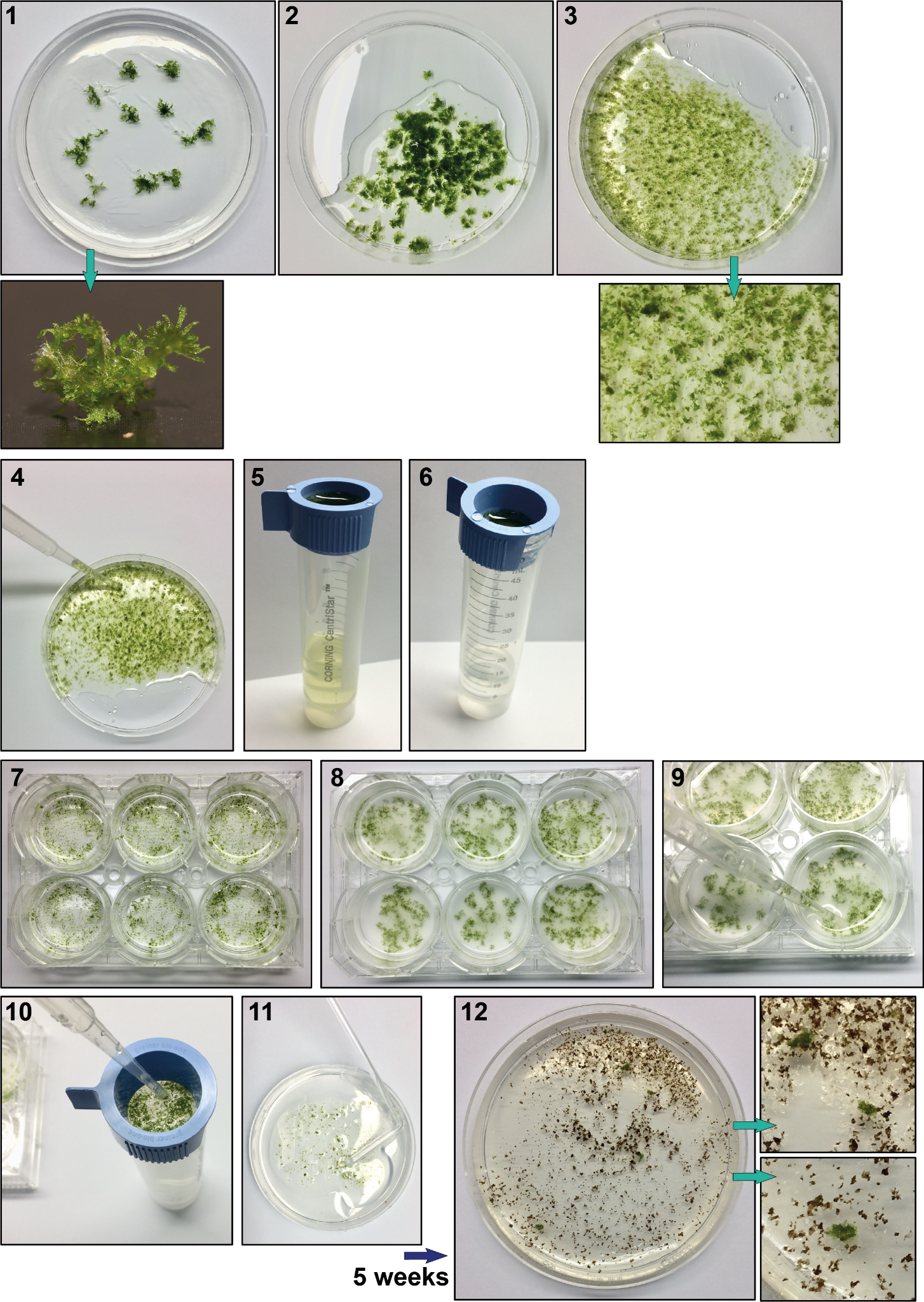

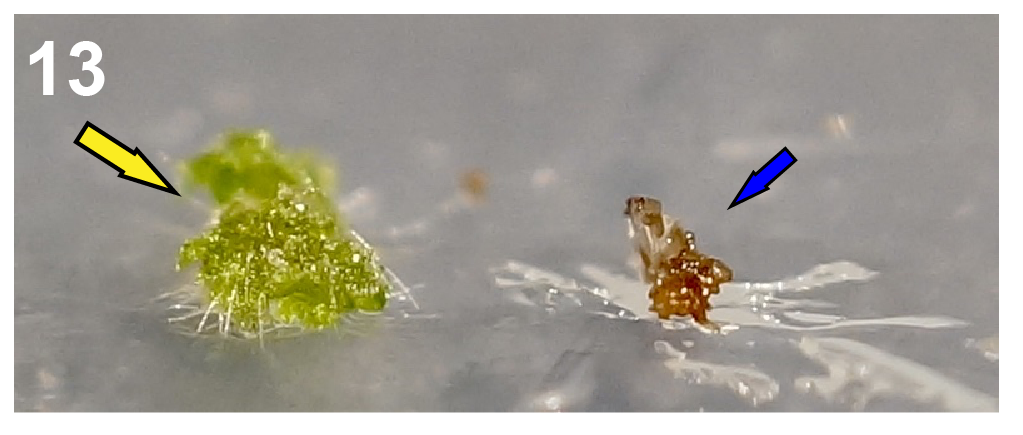

**14**

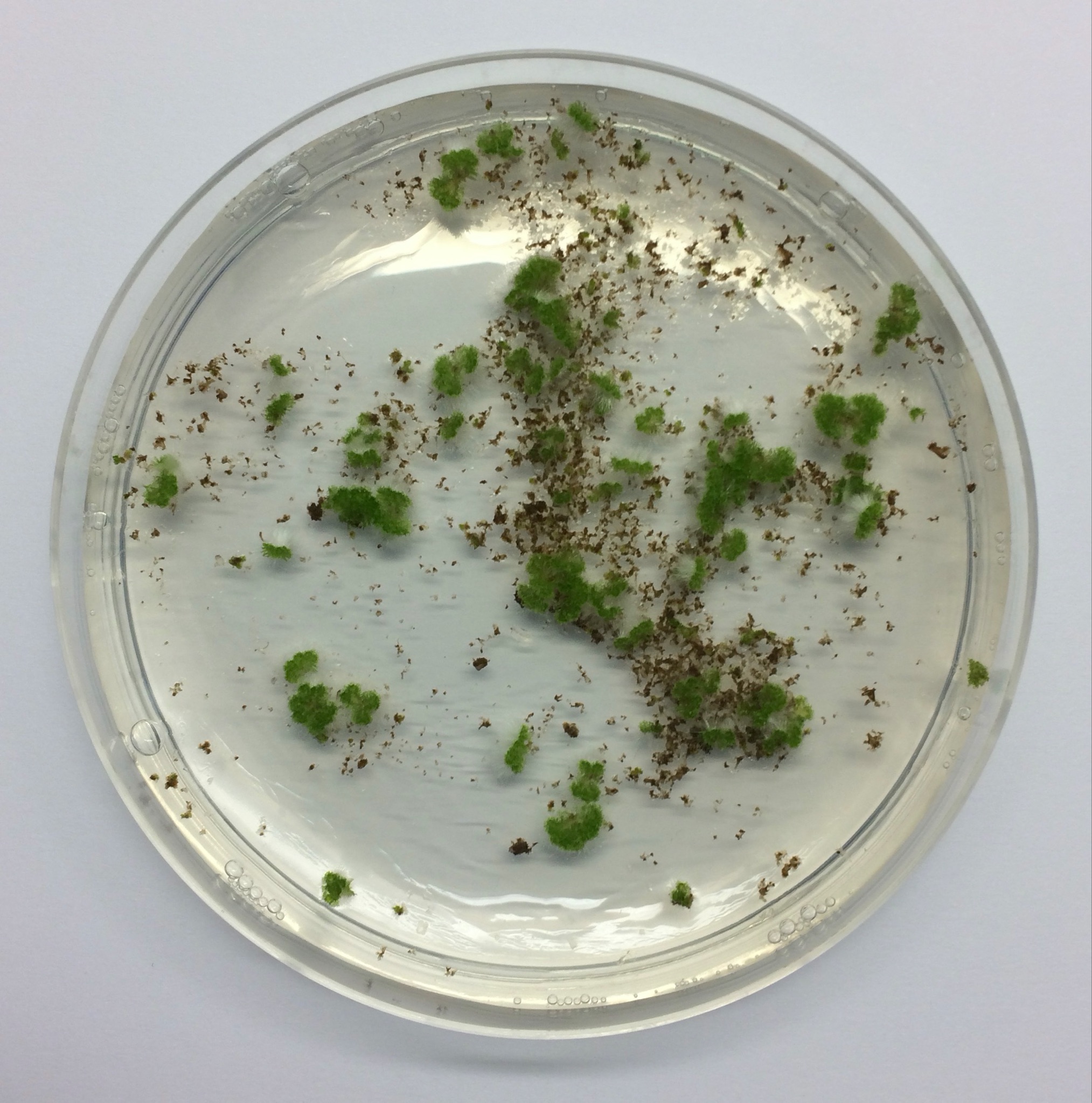

**Figure S5: Workflow showing steps of the transformation protocol optimised for the *A. agrestis* Bonn strain.**

**1)** Approximately 1 g of thallus tissue grown for 4 weeks under low light intensity was collected (approximately 0.1 g of tissue per petri dish - 10 petri dishes in total). **2)** Tissue was transferred into an empty petri dish, sterile water was added until the tissue was covered **3)** the tissue was fragmented using a razor blade (5 mins in total). **4)** the tissue was transferred from the petri dish into a cell strainer positioned on a falcon tube using sterile scalpels. **5-6)** the tissue was washed using ~100 ml of sterile water or until the flow through was clear. **7)** The fragmented thallus tissue was transferred into a 6-well plate (transfer 1⁄6 of the 1 g tissue into a single well) with 5 ml of liquid KNOP medium supplemented with 1% (w/v) sucrose and 40 mM MES, 80 μL of *Agrobacterium* culture and acetosyringone at final concentration of 100 μM. **8)** The tissue was co-cultivated with the *Agrobacterium* for 3 days on a shaker at 110 rpm, with only ambient light. **9-10)** Using a sterile plastic pipette the tissue of one well was transferred into a cell strainer, drained and then transferred on growth media containing the appropriate antibiotic (onto 1 petri dish from one well). **11)** To facilitate spreading of the tissue, 2 ml of sterile water was added to the petri dish. **12)** After 4-6 weeks successful transformants were visible on the petri dish (successful transformants can be identified using a dissecting scope after 4 weeks selection based on rhizoid production and/or fluorescence if such a marker is present on the construct). **13)** The emergence of rhizoids is an indication of successful transformation (yellow arrow: transformed thallus fragment, blue arrow: dying thallus fragment). To eliminate false positives, surviving tissue fragments were transferred again on antibiotics containing growth media. **14)** Example of plate with successful transformants 8 weeks after co-cultivation.

Petri dish dimensions: 92 x16 mm.

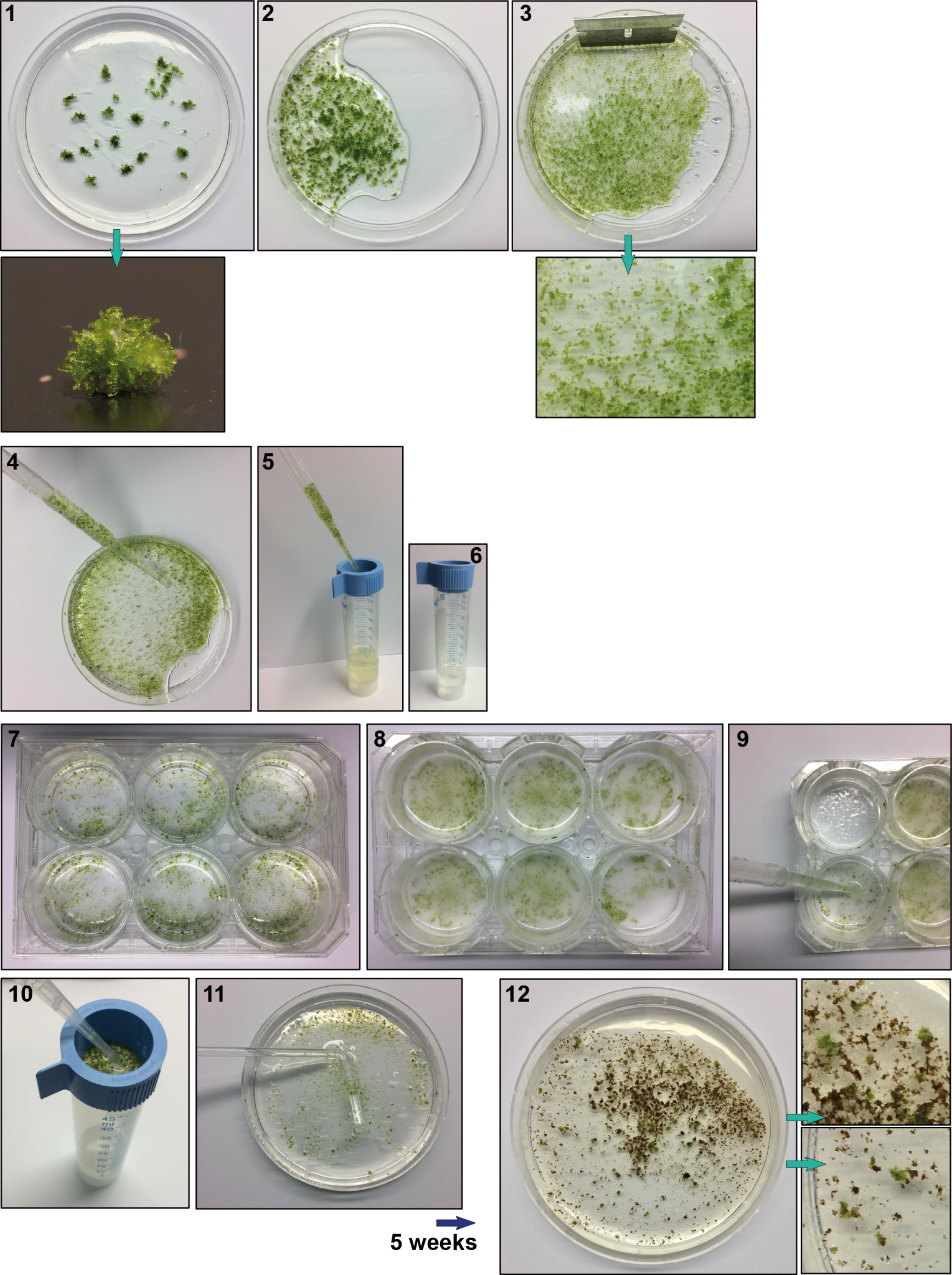

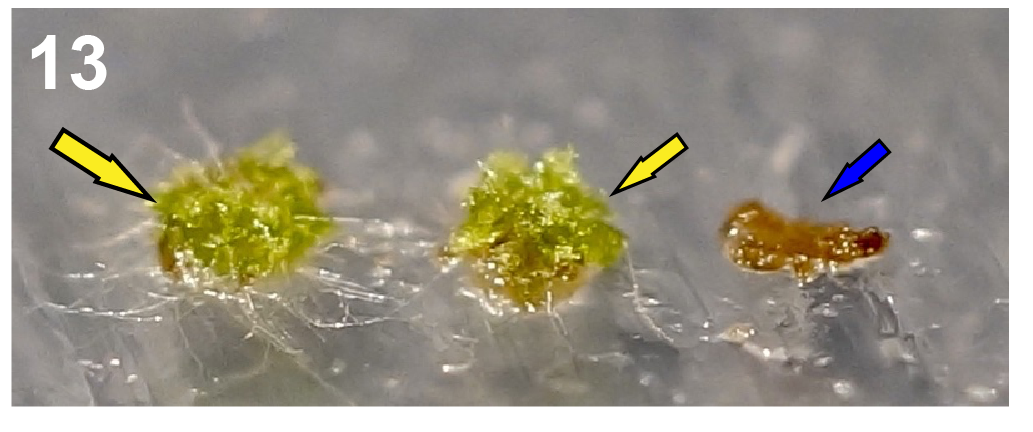

**14**

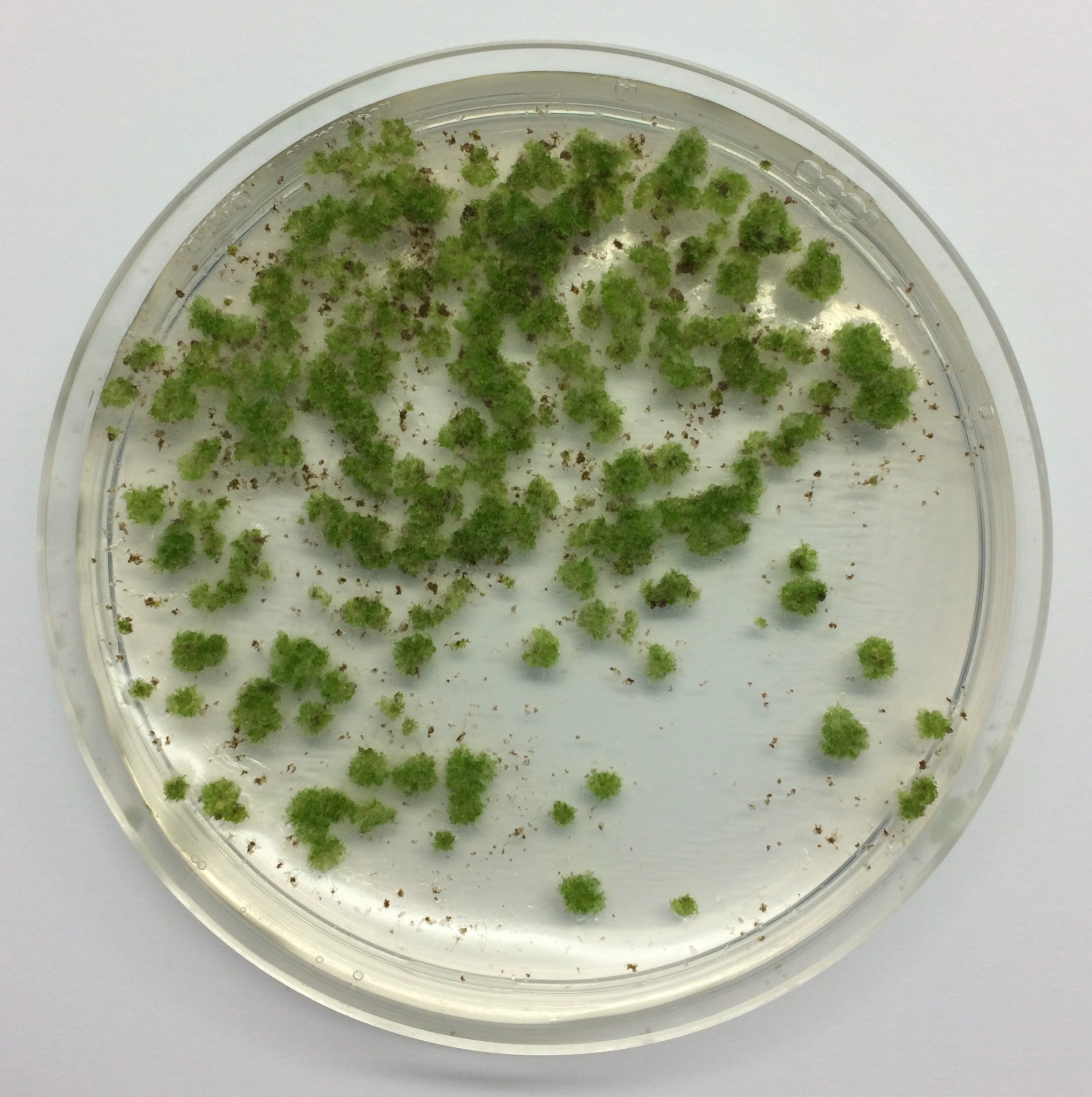

**Figure S6:** **Workflow showing steps of the transformation protocol optimised for the *A. agrestis* Oxford strain.**

**1)** Approximately 1 g of thallus tissue grown for 4 weeks under low light intensity was collected (approximately 0.1 g of tissue per petri dish - 10 petri dishes in total). **2)** Tissue was transferred into an empty petri dish, sterile water was added until the tissue was covered **3)** the tissue was fragmented using a razor blade (5 mins). **4)** the tissue was transferred from the petri dish into a cell strainer positioned on a falcon tube using sterile scalpels. **5-6)** the tissue was washed using ~100 ml of sterile water or until the flow through was clear. **7)** The fragmented thallus tissue was transferred into a 6-well plate (transfer 1⁄6 of the 1 g tissue into a single well) with 5 ml of liquid KNOP medium supplemented with 1% (w/v) sucrose and 40 mM MES, 80 μl of *Agrobacterium* culture and acetosyringone at final concentration of 100 μM. **8)** The tissue was co-cultivated with the *Agrobacterium* for 3 days on a shaker at 110 rpm, with only ambient light. **9-10)** Using a sterile plastic pipette the tissue of one well was transferred into a cell strainer, drained and then transferred on growth media containing the appropriate antibiotic (onto 1 petri dish from one well). **11)** To facilitate spreading of the tissue, 2 ml of sterile water was added to the petri dish. **12)** After 4-6 weeks successful transformants were visible on the petri dish (successful transformants can be identified using a dissecting scope after 4 weeks selection based on rhizoid production and/or fluorescence if such a marker is present on the construct). **13)** The emergence of rhizoids is an indication of successful transformation (yellow arrows: transformed thallus fragment, blue arrow: dying thallus fragment). To eliminate false positives, surviving tissue fragments were transferred again on antibiotics containing growth media. **14)** Example of plate with successful transformants 8 weeks after co-cultivation.

Petri dish dimensions: 92 x16 mm.

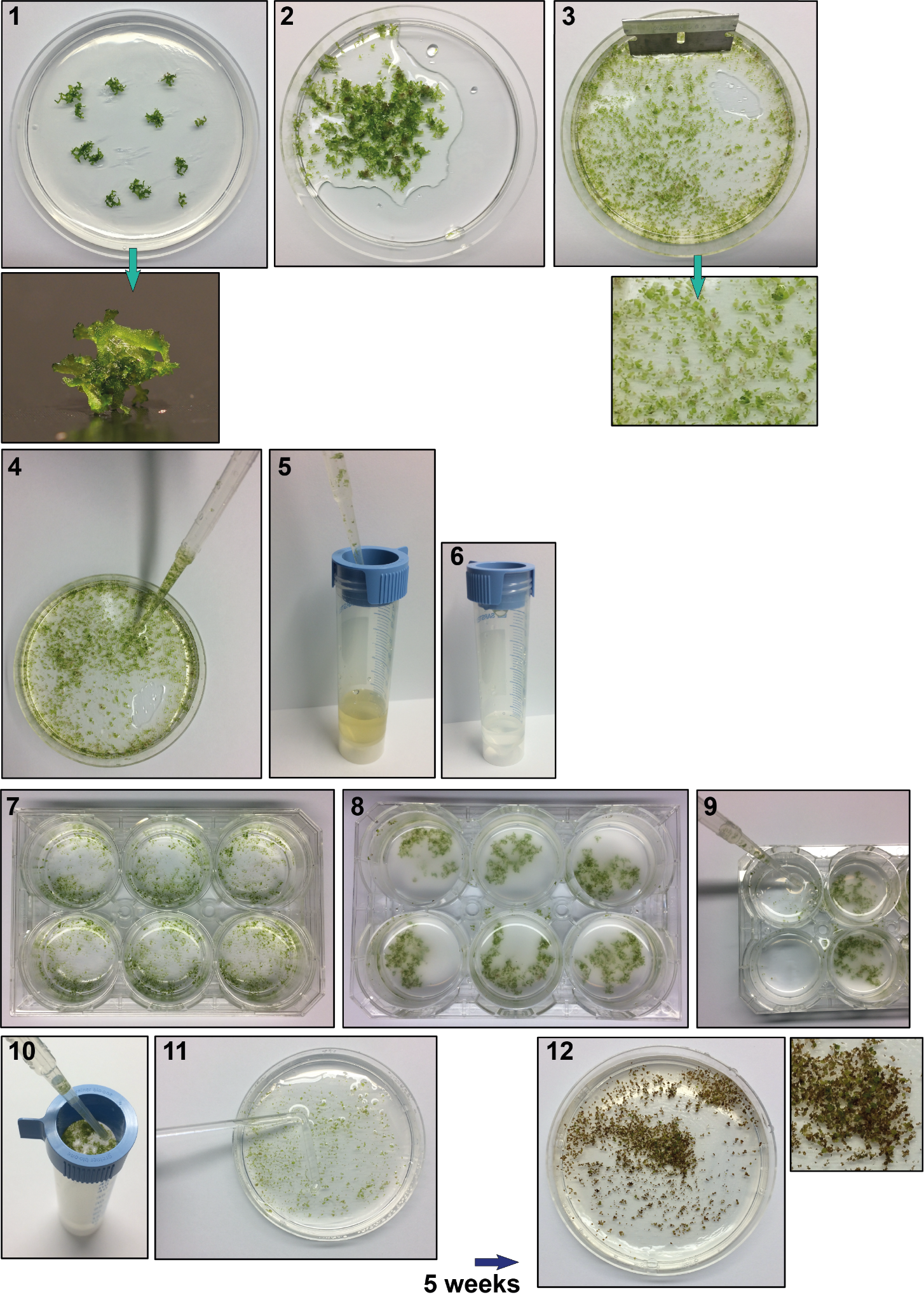

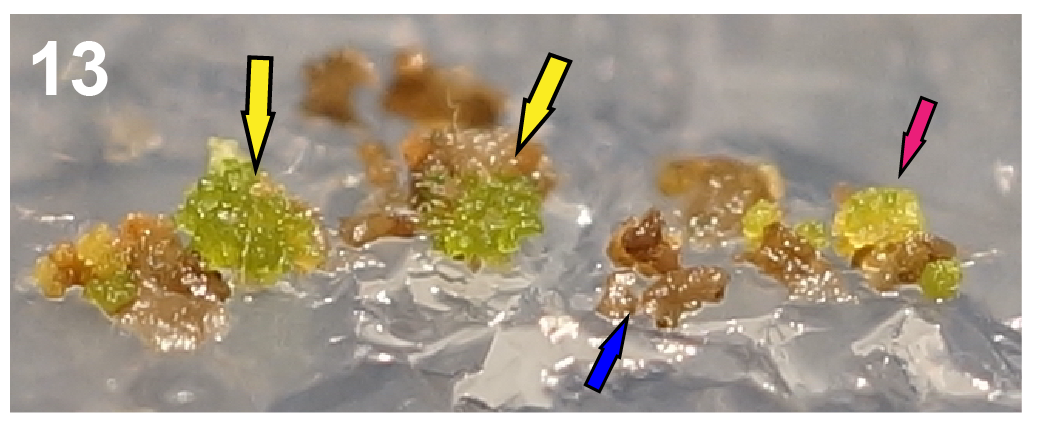

**14**

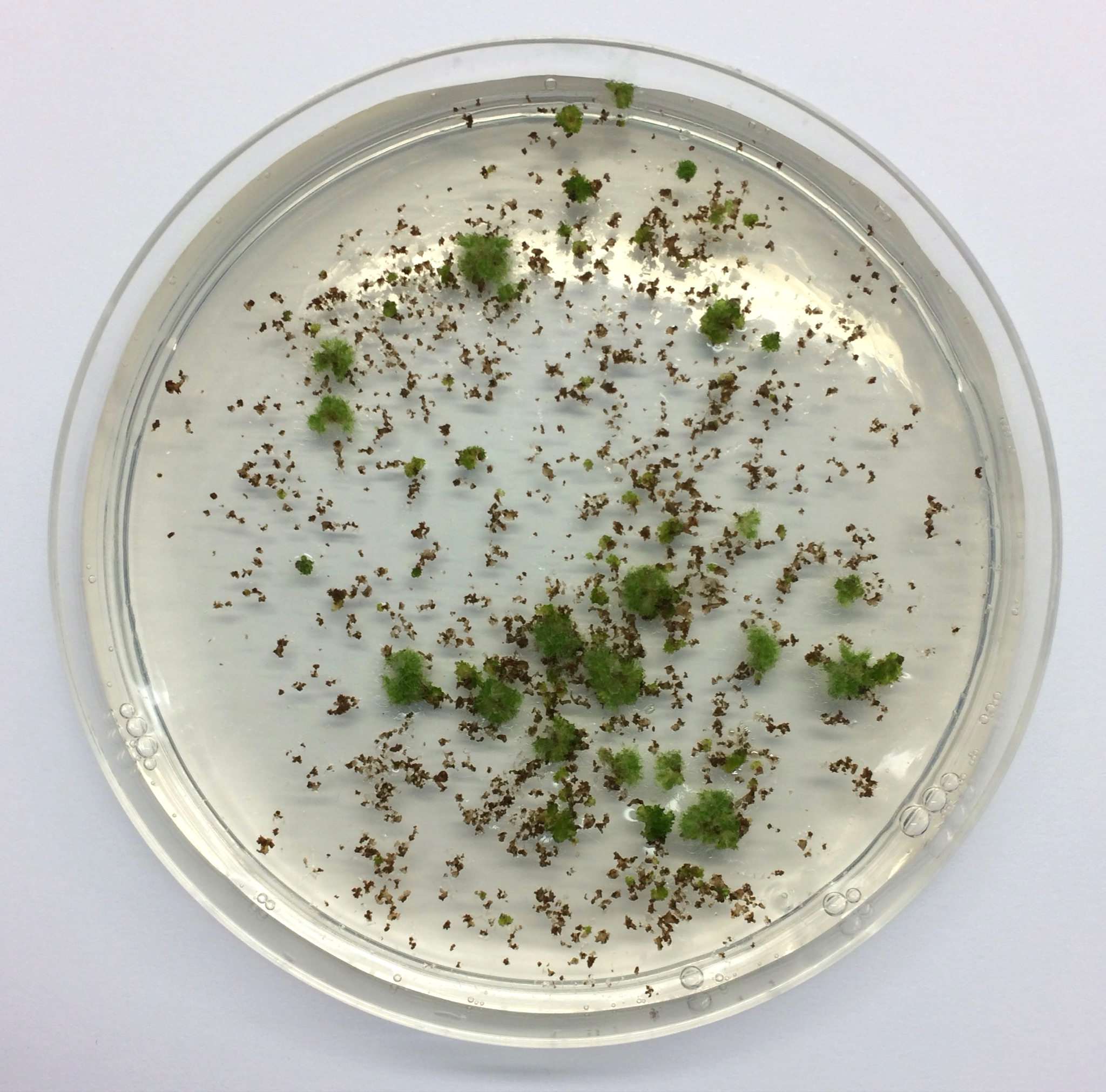

**Figure S7: Workflow showing steps of the protocol used to transform *A. punctatus*.**

**1)** Approximately 1 g of thallus tissue grown for 4 weeks under low light intensity was collected (approximately 0.1 g of tissue per petri dish - 10 petri dishes in total). **2)** Tissue was transferred into an empty petri dish, sterile water was added until the tissue was covered **3)** the tissue was fragmented using a razor blade (5 mins). **4)** the tissue was transferred from the petri dish into a cell strainer positioned on a falcon tube using sterile scalpels. **5-6)** the tissue was washed using ~100 ml of sterile water or until the flow through was clear. . **7)** The fragmented thallus tissue was transferred into a 6-well plate (transfer 1⁄6 of the 1 g tissue into a single well) with 5 ml of liquid KNOP medium supplemented with 1% (w/v) sucrose and 40 mM MES, 80 μL of *Agrobacterium* culture and acetosyringone at final concentration of 100 μM. **8)** The tissue was co-cultivated with the *Agrobacterium* for 3 days on a shaker at 110 rpm, with only ambient light. **9-10)** Using a sterile plastic pipette the tissue of one well was transferred into a cell strainer, drained and then transferred on growth media containing the appropriate antibiotic (onto 1 petri dish from one well). **11)** . To facilitate spreading of the tissue, 2 ml of sterile water was added to the petri dish. **12)** After 4-6 weeks successful transformants were visible on the petri dish (successful transformants can be identified using a microscope after 4 weeks selection based on rhizoid production and/or fluorescence if such a marker is present on the construct). **13)** The emergence of rhizoids is an indication of successful transformation (yellow arrows: transformed thallus fragment, blue and pink arrows: dying thallus fragment).To eliminate false positives, surviving tissue fragments were transferred again on antibiotics containing growth media. **14)** Example of plate with successful transformants 8 weeks after co-cultivation.

Petri dish dimensions: 92 x16 mm.

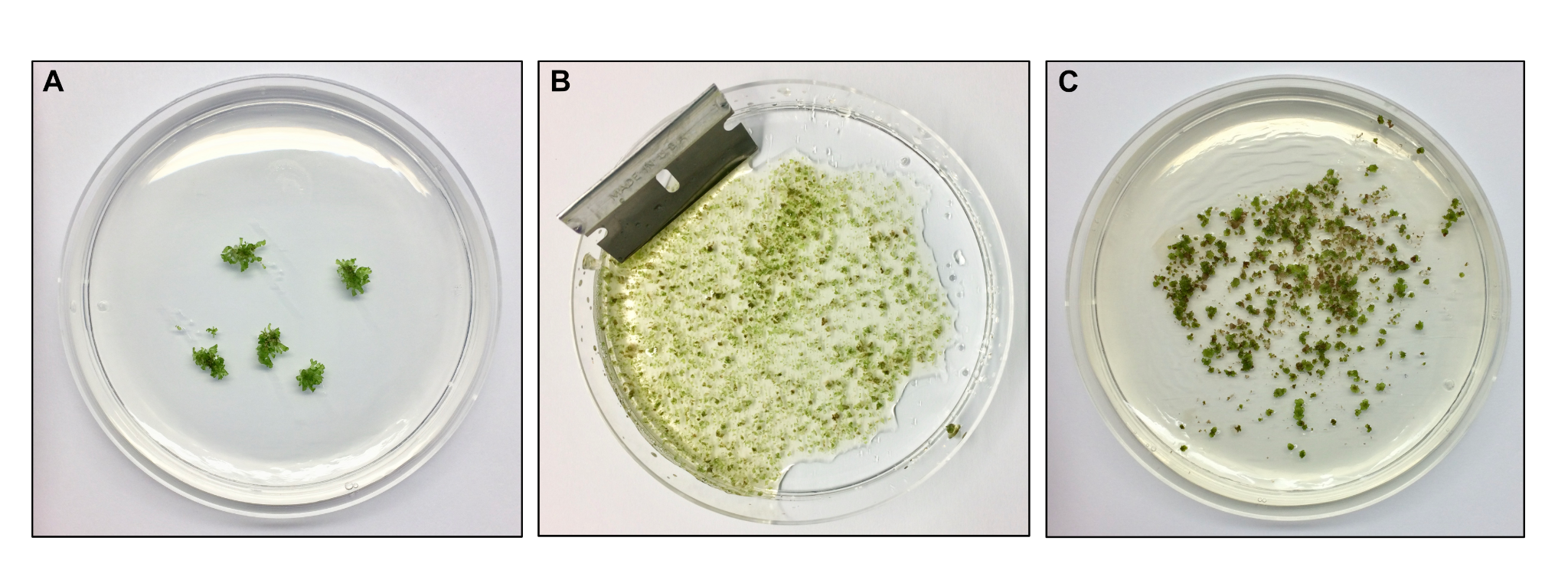

**Figure S8:** **Workflow showing steps of the protocol used to transform *L. dussii***

A) Approximately 1 g of thallus tissue grown for 4 weeks under low light intensity was collected. B) Tissue was transferred into an empty petri dish, sterile water was added until the tissue was covered and then fragmented using a razor blade (5 mins). The following steps are identical to those for the *Anthoceros* species. C) Example of plate with L. dussii on selection 5 weeks after co-cultivation.

Petri dish dimensions: 92 x16 mm.

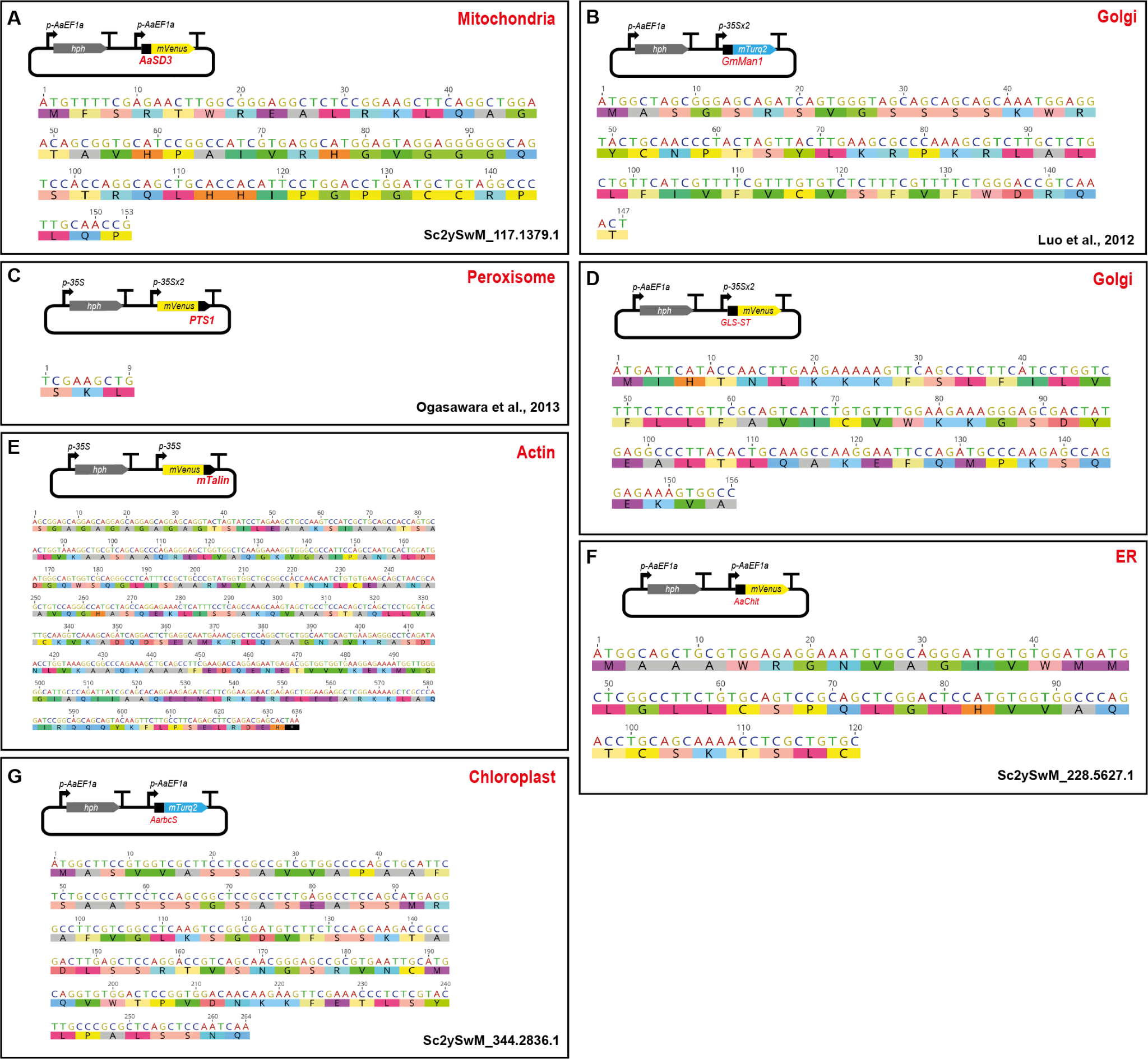

**Figure S9: Summary of sequences used as transit peptides in this study to tailor specific localization of fluorescent proteins.**

Schematic representation of the constructs, amino acid and nucleotide sequences of the targeting peptide tested for A) mitochondria (*p-AaEf1a::hph - AaEF1a::mVenus-AaSD3*), B and C) Golgi (*p-AaEf1a::hph - p-35Sx2::mTurquoise2-GmMan1*) (Luo and Nakata 2012) and (*p-AaEf1a::hph - p-35Sx2::mVenus-GLS-ST*) , D) Peroxisome (*p-35S::hph - p-35Sx2::mVenus-PTS1*) (Ogasawara et al. 2013), E) Actin (*p-35S::hph - p-35S::mVenus-mTalin*), F) ER (*p-AaEf1a::hph - AaEF1a::mVenus-AaChit*) and G) Chloroplast (*p-AaEf1a::hph - AaEF1a::AarbcS-mTurquoise2*). All construct maps at Supp Table 1.

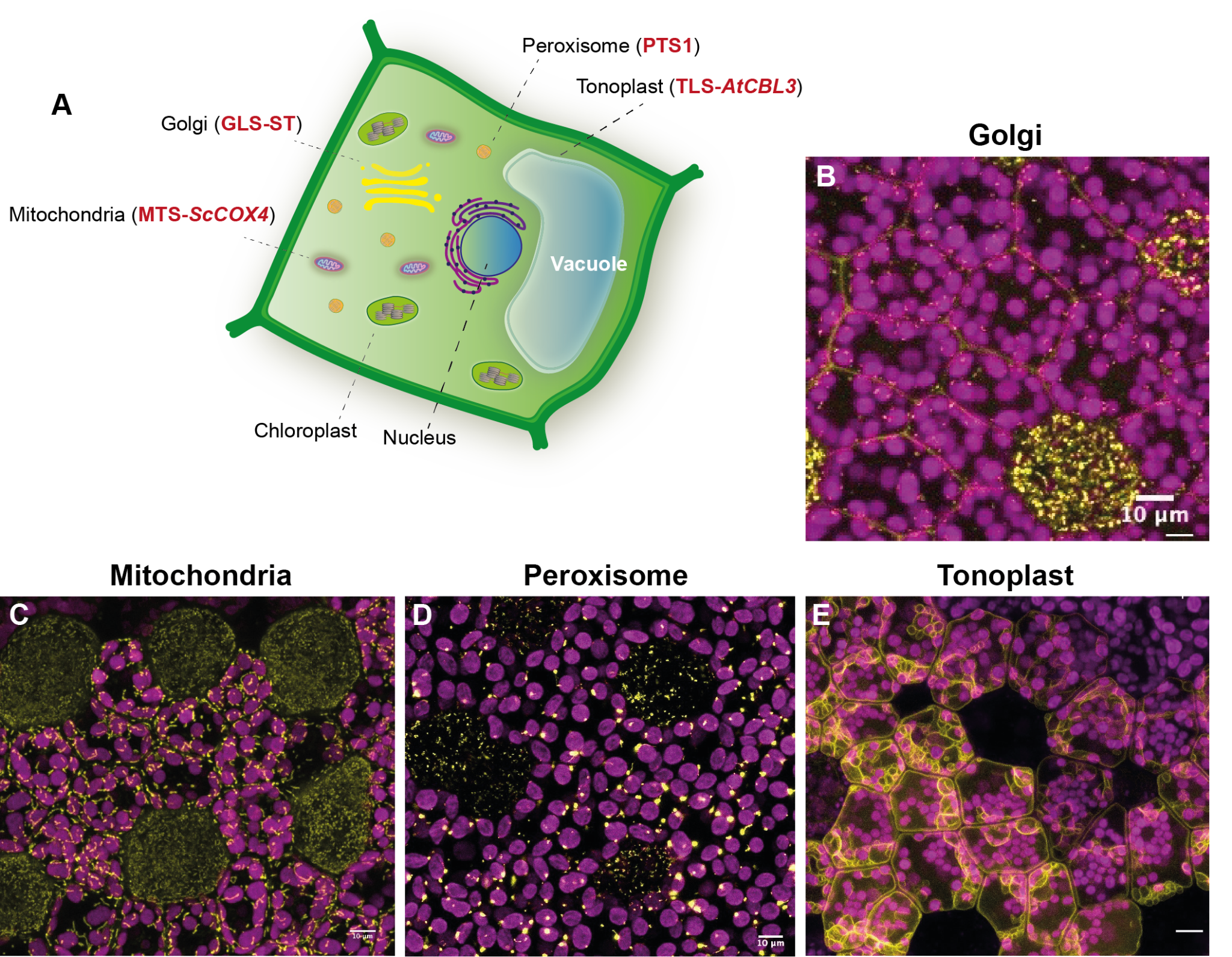

**Figure S10: Localization of fluorescent proteins tagged with various transit peptides in the liverwort *Marchantia polymorpha*.**

A) Schematic representation of a hypothetical *M. polymorpha* cell showing a summary of the subcellular localisation peptides tested in this study. B) Confocal microscopy image of *M. polymorpha* gemmae expressing the Golgi-targeted construct (*p-35S::hph - p-35Sx2::mVenus-GLS-ST*). Scale bar: 10 μm. C) Confocal microscopy image of *M. polymorpha* gemmae expressing the mitochondria-targeted construct (*p-35S::hph - p-35Sx2::mVenus-MTS-ScCOX4*). Scale bar: 10 μm. D) Confocal microscopy image of *M. polymorpha* gemmae expressing the peroxisome-targeted construct (*p-35S::hph - p-35Sx2::mVenus-PTS1*). Scale bar: 10 μm. E) Confocal microscopy image of *M. polymorpha* gemmae expressing the tonoplast-targeted construct (*p-35S::hph - p-35Sx2::mVenus-TLS-AtCBL3*). All construct maps in Supp Table 1.

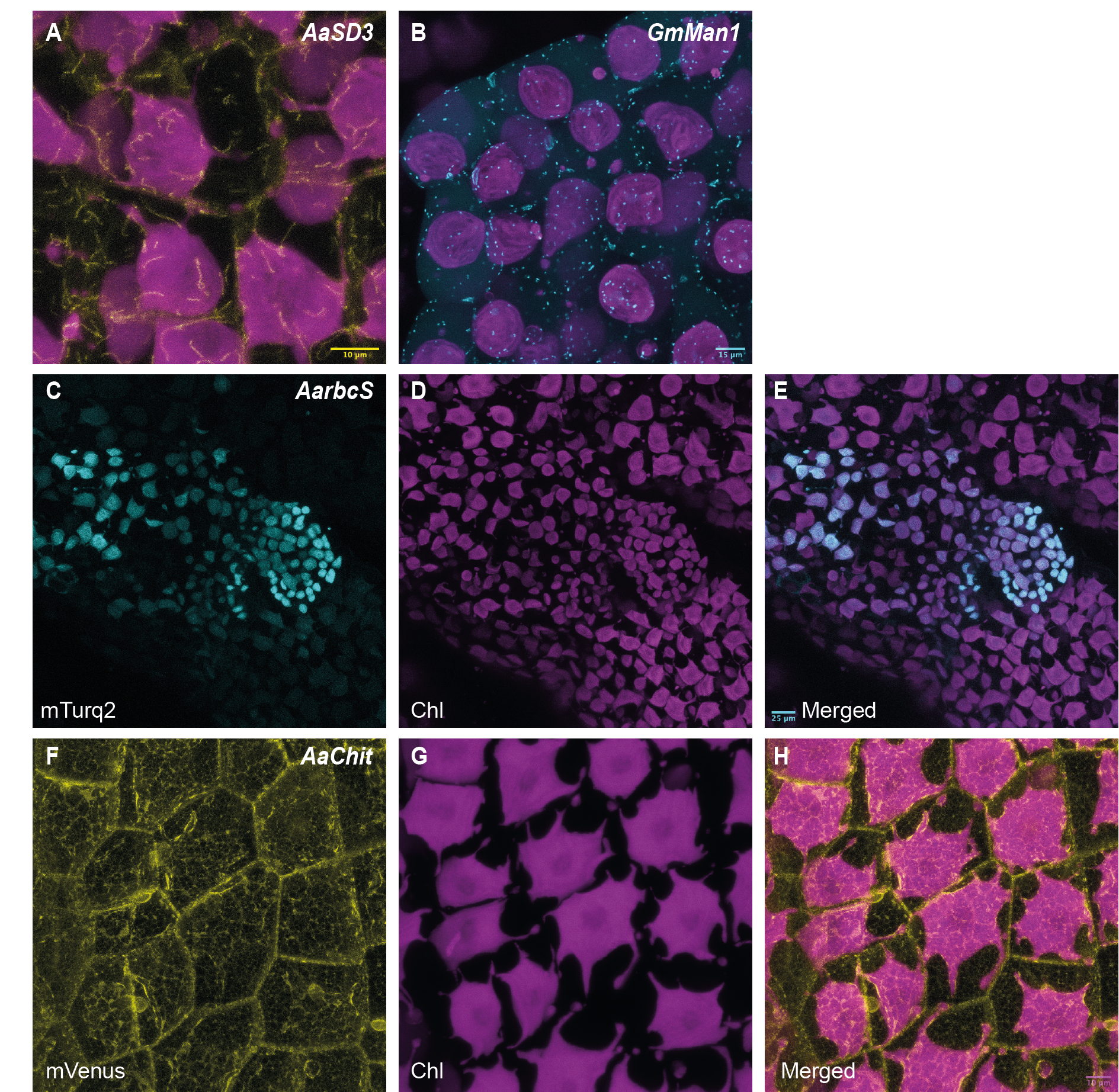

**Figure S11 Targeting fluorescent proteins to various subcellular compartments in the hornwort *A. punctatus***.

A) Confocal microscopy image of *A. punctatus* expressing the *p-AaEf1a::hph - p-AaEF1a::mVenus-AaSD3* construct. Scale bar: 10 μm B) Confocal microscopy image of *A. punctatus* expressing the *p-AaEf1a::hph - p-AaEF1a::mTurquoise-GmMan1* construct. Scale bar: 15 μm. C-E) Confocal microscopy image of *A. punctatus* expressing the *p-AaEf1a::hph - p-AaEF1a::Aa-rbcS-mTurquoise2* construct. Scale bar: 25 μm F-H) Confocal microscopy image of *A. punctatus* expressing the *p-AaEf1a::hph - p-AaEF1a::mVenus-AaChit* construct. Scale bar: 10 μm All construct maps at Supp Table 1.

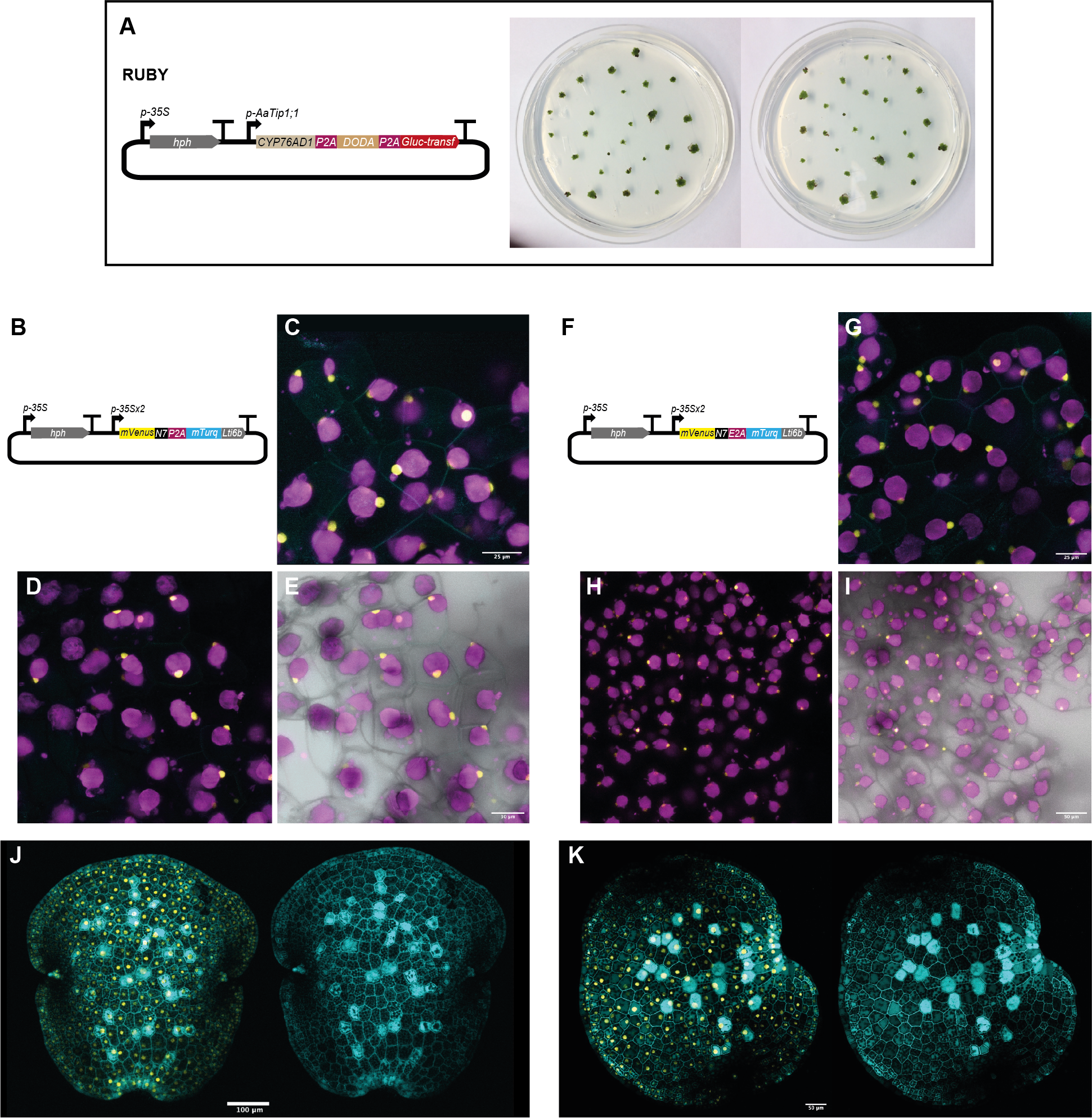

**Figure S12: Testing utility of the RUBY reporter and the 2A self cleavage peptides in *A. agrestis*.**

A) Right: Schematic representation of the RUBY construct. Left: Images of *A. agrestis* Oxford expressing the RUBY construct (*p-35S::hph - p-AaTip1;1::RUBY* - map Suppl Table 1). B) Schematic representation of the construct for P2A self-cleavage peptide testing (*p-35S::hph - p-35Sx2:mVenus-P2A-mTurquoise2-Lti6b*). C-E: Images of *A. agrestis* Oxford expressing the P2A self-cleavage peptide construct. J) Images of *M. polymorpha* expressing the P2A self-cleavage peptide construct. F) Schematic representation of the construct for the E2A self-cleavage peptide testing (*p-35S::hph - p-35Sx2:mVenus-N7-E2A-mTurquoise2-Lti6b*). G-I: Images of *A. agrestis* Oxford expressing the E2A self-cleavage peptide construct. K) Images of *M. polymorpha* expressing the E2A self-cleavage peptide construct. All construct maps at Supp Table 1.

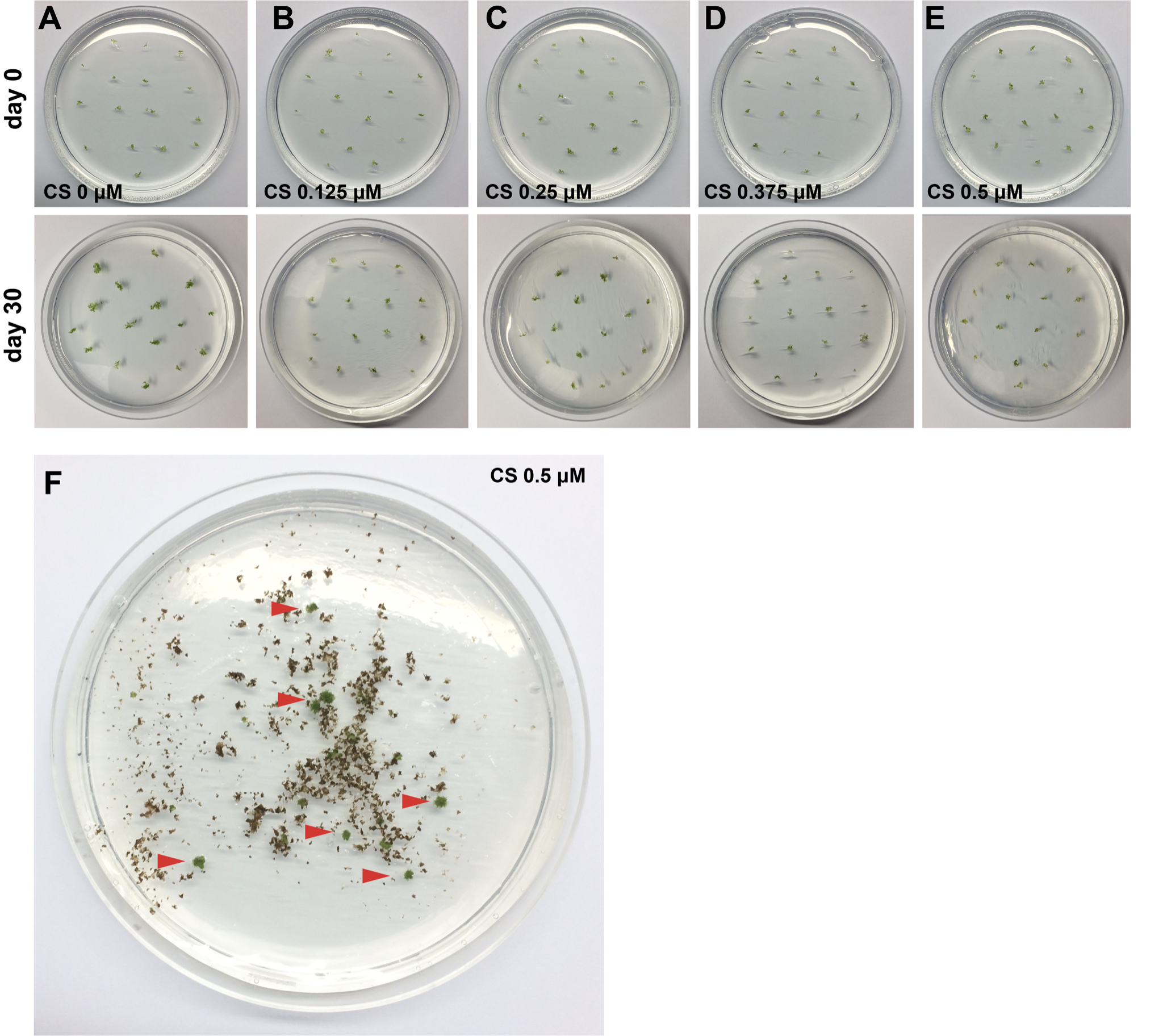

**Figure S13. Chlorsulfuron sensitivity of *A. agrestis* gametophytes.**

A-E) Plates with *A. agrestis* thallus subjected to 0 - 0.5 μM chlorsulfuron selection at day 0 of application of selection (top), day 30 (bottom). F) Example of plate with successful transformants 8 weeks after co-cultivation.

Petri dish dimensions: 92 x16 mm.

**Supplemental video 1: ppt**

**Supplemental Table 1: xls**

Construct sequences (genebank format):

*p-AaEf1a::hph - p-AaTip1;1::eGFP-Lti6b*

*p-AaEf1a::hph - p-AaEF1a::eGFP-Lti6b*

*p-AaEf1a::hph - p-35S_s::eGFP-Lti6b*

*p-AaEf1a::mALS - p-35S_s::eGFP-Lti6b*

*p-35S::hph - p-35Sx2:mVenus-N7-E2A-mTurquoise2-Lti6b*

*p-35S::hph - p-35Sx2:mVenus-P2A-mTurquoise2-Lti6b*

*p-AaEF1a::hph - p-35S::mScarlet-Lti6b*

*p-35S::hph - p-AaTip1;1::RUBY*

*p-AaEf1a::hph - AaEF1a::mVenus-AaSD3*

*p-AaEf1a::hph - p-35Sx2::mTurquoise2-GmMan1*

*p-35S::hph - p-35Sx2::mVenus-PTS1*

*p-35S::hph - p-35S::mVenus-mTalin*

*p-AaEf1a::hph - AaEF1a::AarbcS-mTurquoise2*

*p-AaEf1a::hph - AaEF1a::mVenus-AaChit*

*p-AaEf1a::hph - p-35Sx2::mVenus-GLS-ST*

*p-35S::hph - p-35Sx2::mVenus-TLS-AtCBL3*

*p-35S::hph - p-35Sx2::mVenus-MTS-ScCOX4*

*p-35S::hph - p-35Sx2::mVenus-GLS-ST*

**Supplemental Table 2.**

|  | *construct* | *Species* | *Number of transformants* | *Expression in rhizoids* | *Patchy expression* | *No fluorescence/ red colour* |
| --- | --- | --- | --- | --- | --- | --- |
|  | *p-35S::hph - p-35Sx2:mVenus-N7-E2A-mTurquoise2-Lti6b* | *A. agrestis*  Oxford | *16* | *2* | *14 (only mVenus)* | *1* |
|  | *p-35S::hph - p-35Sx2:mVenus-P2A-mTurquoise2-Lti6b* | *A. agrestis*  Oxford | *19* | *1* | *17 (only mVenus)* | *2* |
|  | *p-35S::hph - p-35Sx2:mVenus-N7-E2A-mTurquoise2-Lti6b* | *A. punctatus* | *7* | *-* | *7 (only mVenus)* | *-* |
|  | *p-35S::hph - p-35Sx2:mVenus-P2A-mTurquoise2-Lti6b* | *A. punctatus* | *5* | *-* | *5 (only mVenus)* | *-* |
|  | *p-35S::hph - p-AaTip1;1::RUBY* | *A. agrestis*  Oxford | *30* | *-* | *-* | *30* |
